# Fc immunoreceptors promote autophagy to regulate monocyte functions

**DOI:** 10.1101/2024.08.29.610296

**Authors:** Mathilde Nugue, Marie D’Allemagne, Despoina Koumantou, Mathias Vetillard, Mark S Cragg, Pierre Bourdely, Sophie Lotersztajn, Loredana Saveanu

**Author notes:** These authors contributed equally to this work (Marie D’Allemagne, Despoina Koumantou).

## Abstract

Receptors for the Fc fragment of immunoglobulin G (FcyRs) are critical in the defense against pathogens and in monoclonal antibody-based therapies. When activated by immune complexes or opsonized particles, FcyRs are endocytosed. Components of the endocytosis machinery are used during autophagy, a process which is triggered by starvation or by activation of specific receptors. In this work, we demonstrate that activation of FcyRs initiates autophagy, characterized by formation of p62 protein puncta and activation of ULK1, a major component of the autophagy initiation complex. Autophagy induction downstream of FcyRs activation involves the protein phosphatase Pp2a and its enzymatic activity, as demonstrated by *in situ* protein labeling. In animal models in which autophagy was inactivated or enhanced in myeloid cells, autophagy negatively regulates pro-inflammatory cytokine production downstream of FcyRs receptors, while being required for FcyRs -mediated antibody-induced cell phagocytosis and myeloid cell survival. Our results suggest that, for antibody-based therapeutic strategies that target the activation of FcyRs, an additional level of control can be obtained by manipulation of autophagy.

## Introduction

Receptors for the Fc fragment of immunoglobulin G (FcyRs) are expressed by a variety of immune cells. Their activation by soluble or cellular immune complexes (ICs) induces a signaling cascade that can lead to complex pro-inflammatory, anti-inflammatory, and immunomodulatory responses that are critical for host protection against pathogens and for the efficacy of antibody-based therapies. In contrast, FcyR signaling can be detrimental in antibody-induced inflammation and autoimmunity. Thus, FcyRs are central to host defense, tissue homeostasis and immunotherapy (Bournazos *et al*, 2016). Humans and mice express several activating FcyRs but only one inhibitory receptor, FcyRIIB in humans and FcyRII in mice. The inhibitory receptor contains an immunoreceptor tyrosine-based inhibitory motif (ITIM) motif in its cytosolic domain, whereas all activating receptors signal through immunoreceptor tyrosine-based activation motifs (ITAM). For most Fc receptors, the ITAM motif is provided by heterodimerization with the Fc receptor y chain encoded by the *FCER1G* gene in humans and *Fcgr1g* in mice (Brandsma *et al*, 2016). The Fc receptor y chain is dispensable for FcyRIIA and C signaling, which contain an ITAM motif in their intracellular tails.

Upon binding of ICs, the FcyRs cluster and their ITAM motifs are phosphorylated by Src kinases (Duchemin *et al*, 1994), such as Lyn, Hck and Fgr, leading to the recruitment of Syk kinase. Once activated, the Syk kinase recruits class I PI3K and PLCy that catalyze the generation of second messengers, such as inositol triphosphate and diacyglycerol, necessary for the Ca^2+^ levels increase and activation of PKC. This early FcyR downstream signaling leads to myeloid cell activation, production of pro-inflammatory cytokines and reactive oxygen species (ROS), phagocytosis of IgG-opsonized particles, antibody-dependent cell cytotoxicity (ADCC) or cell phagocytosis (ADCP). Finally, late signaling pathways, involving activation of the Ras, MEK and MAP kinase families, together with cytokine and chemokine production, increase cell motility, survival and differentiation (Bournazos *et al*, 2016).

The function of FcyRs is closely linked to endocytosis and endocytic trafficking, as FcyRs are internalized together with ICs after activation (Mellman & Plutner, 1984). This internalization is essential for optimal FcyR signaling, because similar to other ITAM-bearing receptors such as the B cell receptor (BCR) (Chaturvedi *et al*, 2011) and T cell receptor (TCR) (Evnouchidou *et al*, 2020), FcyRs use endosomal signaling platforms whose destabilization compromises the pro-inflammatory response and the ADCC mediated by these receptors (Bratti *et al*, 2022; Benadda *et al*, 2023).

An important pathway that strongly communicates with the endocytic components is autophagy, a process of cellular catabolism triggered by nutrient and energy deprivation. Canonical autophagy can be either bulk, which implies non-selective degradation of cellular components that is triggered by starvation, or selective, which removes damaged organelles or non-functional proteins (Hurley & Young, 2017). Canonical autophagy is initiated by activation of the ULK1 (Atg1) complex, which comprises the kinase ULK1 and the non-catalytic subunits FIP200, ATG13 and ATG101 (Ganley *et al*, 2009). The phosphorylation status of ULK1 is the key determinant for autophagy induction. Under normal nutrient conditions, ULK1 is phosphorylated at Ser757 and Ser637, which maintains ULK1 in an inactive conformation and prevents autophagy induction. Of these two residues, Ser757 is phosphorylated by mTor1, while Ser637 is targeted by several kinases and phosphatases, representing an important site for regulating the induction of autophagy pathway. mTor1 and AMPK1 can both phosphorylate Ser637 to keep ULK1 inactive, while the phosphatases PP2A and PPM1D can dephosphorylate it to induce autophagy (Torii *et al*, 2016; Wong *et al*, 2015). Once activated, ULK1 undergoes autophosphorylation to enhance its activation (Lazarus *et al*, 2015). Autophagosome elongation, closure, and maturation are driven by the ubiquitin-like conjugation systems including ATG3, ATG4, ATG5, ATG7, ATG10, and ATG12 and the lipidation of LC3 (Atg8) (Hurley & Young, 2017). In addition to canonical autophagy, alternative forms of autophagy are LC3-associated phagocytosis (LAP) and LC3-associated endocytosis (LANDO), in which LC3 is recruited to the phagosomal or endosomal single membrane upon activation of innate immune receptors (Sanjuan *et al*, 2007; Heckmann *et al*, 2019). Although LAP/LANDO requires many ATG proteins, they do not require the ULK1 complex, but rather the protein Rubicon. Rubicon promotes LAP and LANDO (Heckmann *et al*, 2019), while blocking canonical autophagy through inhibitory interactions with the class III PI3K subunit Vps34 and with UVRAG, a protein required for autophagosome maturation (Minami *et al*, 2021). In addition, by blocking Rab7 activity, Rubicon inhibits endosome maturation (Sun *et al*, 2010).

In summary, autophagy is a complex pathway that closely interacts with endocytic trafficking (Hurley & Young, 2017), an essential player in FcyR signaling and function (Benadda *et al*, 2023; Bratti *et al*, 2022). Therefore, we wondered whether there is a crosstalk between autophagy and FcyR function in myeloid cells. We show that FcyR activation triggers autophagy via the phosphatase Pp2a and that autophagy is essential for the control of FcyR-mediated functions in myeloid cells. Following activation of FcyRs, autophagy limits pro-inflammatory cytokine production and is required for antibody dependent cell phagocytosis (ADCP), as well as for myeloid cell fitness.

## Results

### FcyRs trigger autophagy via the phosphatase Pp2a

Since endocytosis is involved in FcyRs signaling (Benadda *et al*, 2023; Bratti *et al*, 2022) and the autophagy pathway is tightly linked to endocytosis (Yamamoto *et al*, 2023), we asked whether FcyR activation could activate the autophagy pathway. To this end, we activated FcyRs with ICs and monitored autophagy levels. First, we analyzed the kinetics of induction of p62 puncta, a receptor that captures cytosolic proteins for their integration into the autophagosome. As shown in Figure 1A-B, the number of p62 puncta per cell was increased 30 minutes after FcyR activation, suggesting the activation of autophagy. At later time points, the number of p62 puncta decreased, as expected, since p62 is degraded in autophagolysosomes during autophagosome maturation (Bjørkøy *et al*, 2009). We verified that blocking autophagy flux with chloroquine (CQ) led to a further increase in the number of p62 puncta (Figure 1C-D). Analysis of LC3 distribution by confocal microscopy demonstrated an increase of LC3 positive vesicles after cell incubation with IC, suggesting also the induction of autophagy downstream FcyR activation (Figure 1E-F). To confirm that FcyR activation triggers autophagy, we analyzed the phosphorylation status of ULK1, a key kinase involved in autophagy induction (Chan *et al*, 2007). As described above, ULK1 harbors several inhibitory phosphorylation sites, such as the Ser757 and Ser638 residues, which are phosphorylated by the mTorc1 complex (Wong *et al*, 2013). As shown in Figure 1G-H, the inhibitory phosphorylation of ULK 1 (Ser757) was reduced upon FcyR activation to a level similar to that observed upon treatment with rapamycin, an mTor inhibitor (Figure 1G-H, left panel). As a consequence of rapamycin treatment, the mTorc pathway was inhibited, as shown by the reduction of the active form of S6K70 (pThr389) (Figure 1G-H, right panel), a downstream mTorc effector (Liu & Sabatini, 2020). Taken together, these results demonstrate a significant induction of the autophagy pathway upon activation of FcyRs.

**FIGURE 1:**
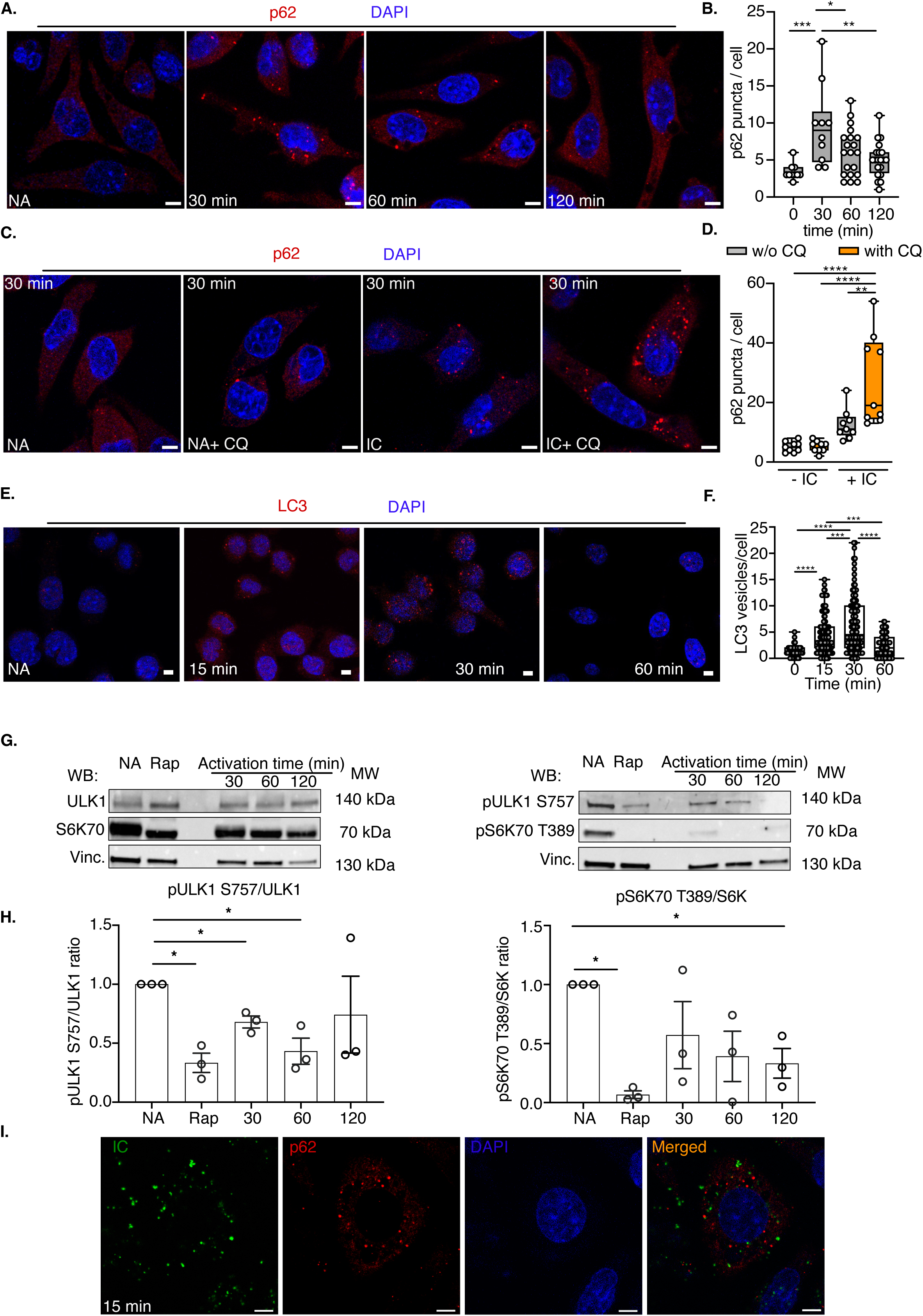
FcyR activation triggers autophagy. (**A-B**) RAW cells were incubated without (NA) or with mouse IgG and anti-mouse IgG at 4°C. After removal of excess antibodies, cells were incubated for the indicated times at 37°C, fixed and stained for p62 (red). The pictures show representative images from 3 independent experiments and the graph shows the number of p62 puncta per cell at the indicated time point. Each dot represents one cell. Scale bars = 5 μm. (**C-D**) RAW cells were incubated without (NA) or with mouse IgG and anti-mouse IgG at 4°C. After removal of excess antibodies, cells were incubated for 30 minutes at 37°C, with or without chloroquine treatment (CQ), fixed and stained for p62 (red). The pictures show representative images from 3 independent experiments and the graph shows the number of p62 puncta per cell at the indicated time point. Each dot represents one cell. Scale bars = 5 μm. (**E-F**) RAW cells were incubated without (NA) or with mouse IgG and anti-mouse IgG at 4°C. After removal of excess antibodies, cells were incubated for the indicated times at 37°C, fixed and stained for LC3 (red). The pictures show representative images from 2 independent experiments and the graph shows the number of p62 puncta per cell at the indicated time points. Each dot represents one cell. Scale bars = 5 μm. (**G-H**) Bone-marrow derived macrophages (BMDMs) were incubated without (NA) or with mouse IgG and anti-mouse IgG at 4°C. After removal of excess antibodies, the cells were were incubated for the indicated times at 37°C. Cells treated with rapamycin (Rap) were incubated at 37°C for 30 minutes. The cells were lysed and the cleared lysate was analyzed by immunoblotting for total ULK1, S6K70 (left panel) and the inhibitory phosphorylation of ULK1 (S757) and activatory phosphorylation of S6K70 (T389). Vinculin (Vinc.) was used as a loading control. The graph in the panel **H** shows the quantification of 3 immunoblot experiments performed as in **G**. (**I**) RAW cells were incubated with mouse IgG and anti-mouse IgG conjugated to Alexa-Fluor647 (green) at 4°C. After removal of excess antibody, cells were incubated at 37°C for 15 minutes, fixed and stained for p62 (red). The pictures show representative images from 2 independent experiments. Scale bars= 5 µm.

It has been previously shown that FcyRs can be internalized by LC3-associated phagocytosis (LAP) and then induce anti-inflammatory signaling from the LAPosomes (Wan *et al*, 2020). However, as shown by the ULK1 phosphorylation experiments (Figure 1G-H), FcyR cross-linking by IC induced canonical autophagy and not LAP.

We did not observe a colocalization between IC and the p62 puncta, indicating that FcyR signaling platforms do not assemble at the autophagosome level (Figure 1I). Thus, FcyR signaling itself seems to induce canonical autophagy by an unknown mechanism. To investigate the involved mechanisms, we performed an *in situ* biotinylation assay using a chimeric protein obtained by fusing FcyRIIA to the APEX2 peroxidase, which biotinylates proteins in a radius of 20 nm (Lobingier *et al*, 2017). The FcyRIIA-GFP-APEX2 fusion protein was expressed in DC2.4 cells and the cells were activated with mouse F(ab’)_2_ anti-FcyRIIA, followed by receptor aggregation with a goat F(ab’)_2_ anti-mouse F(ab’)_2_. The biotinylated proteins were purified with streptavidin-coupled beads and the purified material was immunoblotted for proteins involved in autophagy regulation.

As expected, the major FcyR signaling kinase, Syk, was found close to the receptor only after FcyRIIA activation (Figure 2A). In contrast, autophagy regulatory proteins (mTor, AMPKα a and Pp2a) were already present in proximity of the FcyRIIA and further enriched after its activation (Figure 2B). Since autophagy is known to be regulated by complex mechanisms of phosphorylation and dephosphorylation, we focused on the phosphatase Pp2a, which has been shown to dephosphorylate the key kinase ULK1 at Ser637, leading to ULK1 activation and autophagy induction (Wong *et al*, 2015).

**FIGURE 2:**
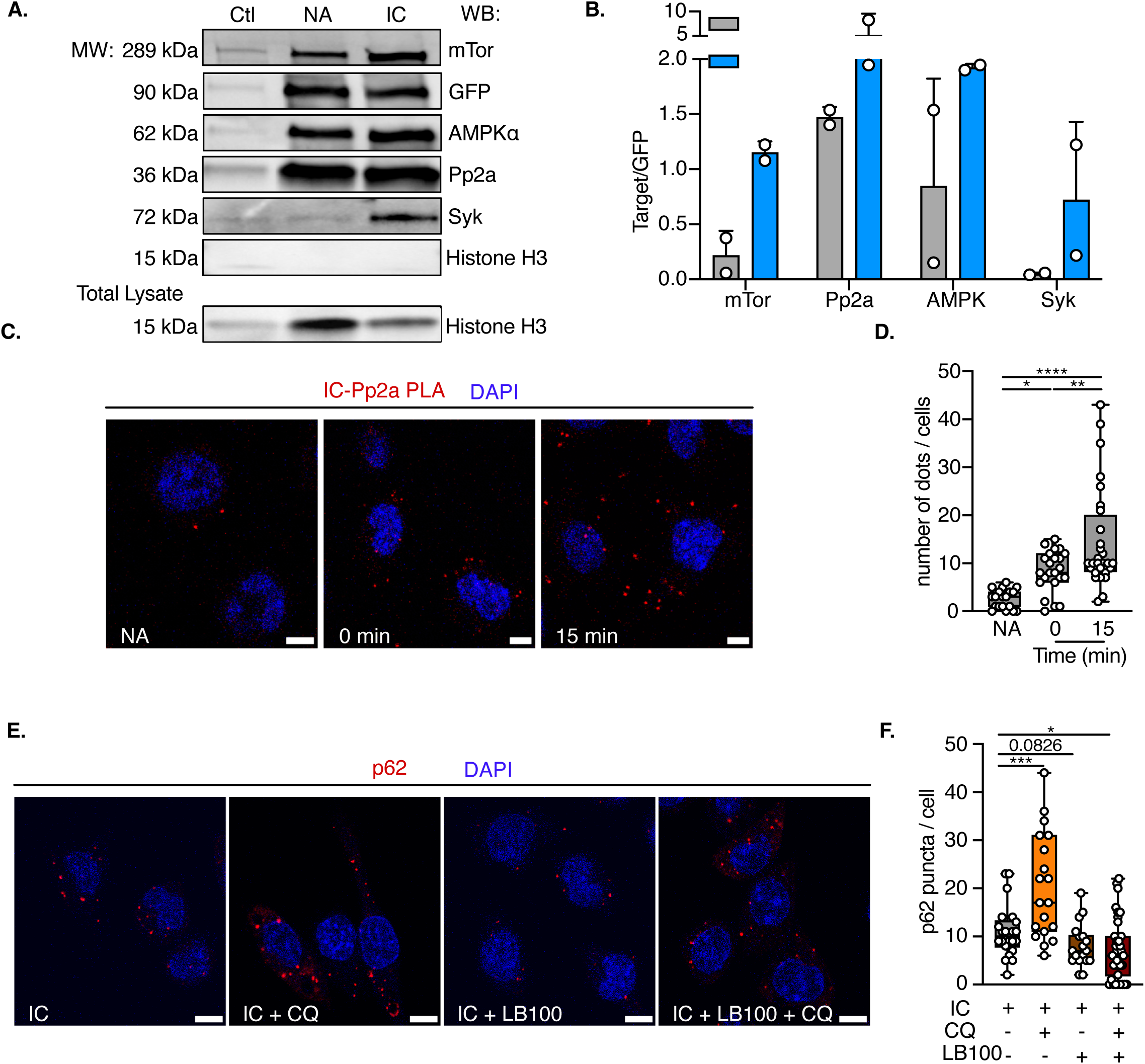
FcyRs recruit the phosphatase Pp2a to induce autophagy. (**A-B**) DC2.4-FcyRIIA-GFP-APEX2 cells were incubated with or without biotin-phenol for 30 minutes prior to activation (or not) with F(ab)’_2_ anti-FcyRIIA and F(ab)’_2_ anti-mouse F(ab)’_2_ at 4°C. After removal of excess antibody, cells were incubated at 37°C for 30 minutes. H_2_O_2_ was added for 30 seconds to allow biotinylation. Cells were then lysed, and biotinylated proteins were recovered using streptavidin magnetic beads. Bead eluates were analyzed by immunoblotting using antibodies specific for Syk, mTor, ULK1, GFP, Pp2a, AMPKa and histone H3. The graph represents quantitative analysis of the ratio target to GFP. The pictures show representatives images from 2 independent experiments. (**C-D**) RAW cells were incubated without (NA) or with mouse IgG and anti-mouse IgG at 4°C. After removal of excess antibody, cells were incubated for 30 minutes at 37°C, fixed, and stained for PP2A. ICs and PP2A were then stained with anti-mouse minus and anti-rabbit plus probes, followed by hybridization, ligation, and amplification steps. Red dots represent the interaction site between ICs and PP2A. The pictures show representative images from 2 independent experiments, and the graphs show the number of red dots per cell at each time point. Each dot represents one cell. Scale bars = 5 μm (**E-F**) Raw cells were incubated with PP2A inhibitor (LB100) 2 hours before receptor activation with mouse IgG and anti-mouse IgG at 4°C. After removal of excess antibodies, cells were incubated for 15 minutes at 37°C, with or without CQ and LB100, fixed and stained for p62 (red). The pictures show representative images from 1 experiment, and the graphs show the number of p62 puncta per cell for each condition. Each dot represents one cell. Scale bars = 5 μm.

To validate the interaction between the phosphatase Pp2a and the IC, we performed a proximity-ligation assay (PLA), which detects protein-protein interaction within a 40 nm radius. Similar to the *in situ* biotinylation assay (Figure 2A-B), the PLA assay using anti-IC and anti-Pp2a antibodies detected an interaction between IC and Pp2a as soon as the ICs were added to the cells (time 0 min) and this interaction was significantly increased at 15 min after FcyR activation (Figure 2C-D). To investigate whether the enzymatic activity of Pp2a was required for the induction of the autophagy upon FcyR activation by IC, we repeated the p62 puncta assay upon FcyR activation in the presence of LB100, a Pp2a inhibitor (Hong *et al*, 2015). In the presence of LB100, the number of p62 puncta per cell was reduced at 15 minutes after FcyR activation, as compared with untreated cells. This reduction was more pronounced in the presence of chloroquine, which blocks autophagic cargo degradation and thus p62 puncta degradation (Figure 2E-F).

Taken together, these results indicate that FcyR activation by ICs triggers the autophagy pathway through Pp2a, whose enzymatic activity is essential for the subsequent induction of autophagy. Despite autophagy activation, autophagosomes do not appear to be a signaling platform for FcyRs. However, modulation of autophagy itself could affect FcyR expression, trafficking, signaling and function in phagocytic cells.

### Autophagy is required for FcyR-mediated cytokine production and antibody-dependent cell phagocytosis

To investigate whether autophagy controls FcyR function, we generated mice in which the autophagy pathway was negatively or positively modulated in myeloid cells. These mouse strains were generated by crossing *Lyz2*-Cre mice with *Atg5*^loxlox^ mice (Atg5 -/-) or *Rubicon*^loxlox^ mice (Rubcn -/-) (Figure 3A-B). Atg5 is an essential protein for autophagy initiation and progression (Yamamoto *et al*, 2023), while Rubicon is an autophagy inhibitor which sequesters UVRAG and Rab7 (Matsunaga *et al*, 2009; Zhong *et al*, 2009). Canonical autophagy is known to regulate inflammatory responses downstream of several immune receptors that recognize pathogen-associated molecular patterns (PAMPs), danger associated molecular patterns (DAMPs) and cytokines (Wu & Lu, 2019), but, to our knowledge, its role in pro-inflammatory responses downstream of FcyR activation is unknown. To investigate the role of autophagy in cytokine production downstream of FcyR activation, we activated Atg5 -/- and Rubcn -/- bone marrow derived macrophages (BMDM) with IC alone or in combination with the TLR4 ligand, LPS. Addition of ICs alone to wild-type macrophages did not induce the production of mRNA coding for pro-inflammatory cytokines, whereas in the same situation Atg5 -/- macrophages produced significant levels of IL-1β and TNF-α mRNA. Treatment with LPS alone or in combination with ICs led to higher levels of IL-1β and TNF-α in Atg5-deficient macrophages than in wild-type, suggesting that autophagy exerts a negative control of the FcyR- and TLR4-induced inflammatory response (Figure 3C, left panel). In contrast, activation of Rubcn -/- macrophages by ICs did not exhibit elevated IL-1β or TNF-α production. When TLR4 was stimulated, the absence of Rubicon resulted in increased IL-1β production and a sharp decrease in TNF-α. This suggests that, similar to other immune receptors, autophagy regulates cytokine production at the transcriptional level downstream of FcyR activation. To investigate the pathophysiological relevance of our findings, we exposed Atg5-/- and Rubcn-/- to anti-collagen antibodies and LPS, a model of FcyR-mediated arthritis (Khachigian, 2006). Despite modulation of cytokine production by autophagy-modified macrophages *in vitro*, neither Atg5, nor Rubicon deficiency affected the arthritis score *in vivo* (Supplementary Figure 1). These results may be explained by the fact that Lys2 is highly expressed in osteoclasts, which play a key role in the development of arthritis. For example, ATG7-deficiency in myeloid cells has been shown to reduce osteoclast differentiation and protect the mice against bone erosion (Lin *et al*, 2013). The lack of any effect of Rubicon deletion on the progression of arthritis can be explained by reduced production of reactive oxygen species (ROS), since the chemical disruption of the interaction between Rubicon and the NADPH oxidase NOX2 or Rubicon deficiency reduces ROS production by myeloid cells and protects the mice against arthritis (Kim *et al*, 2020).

**FIGURE 3:**
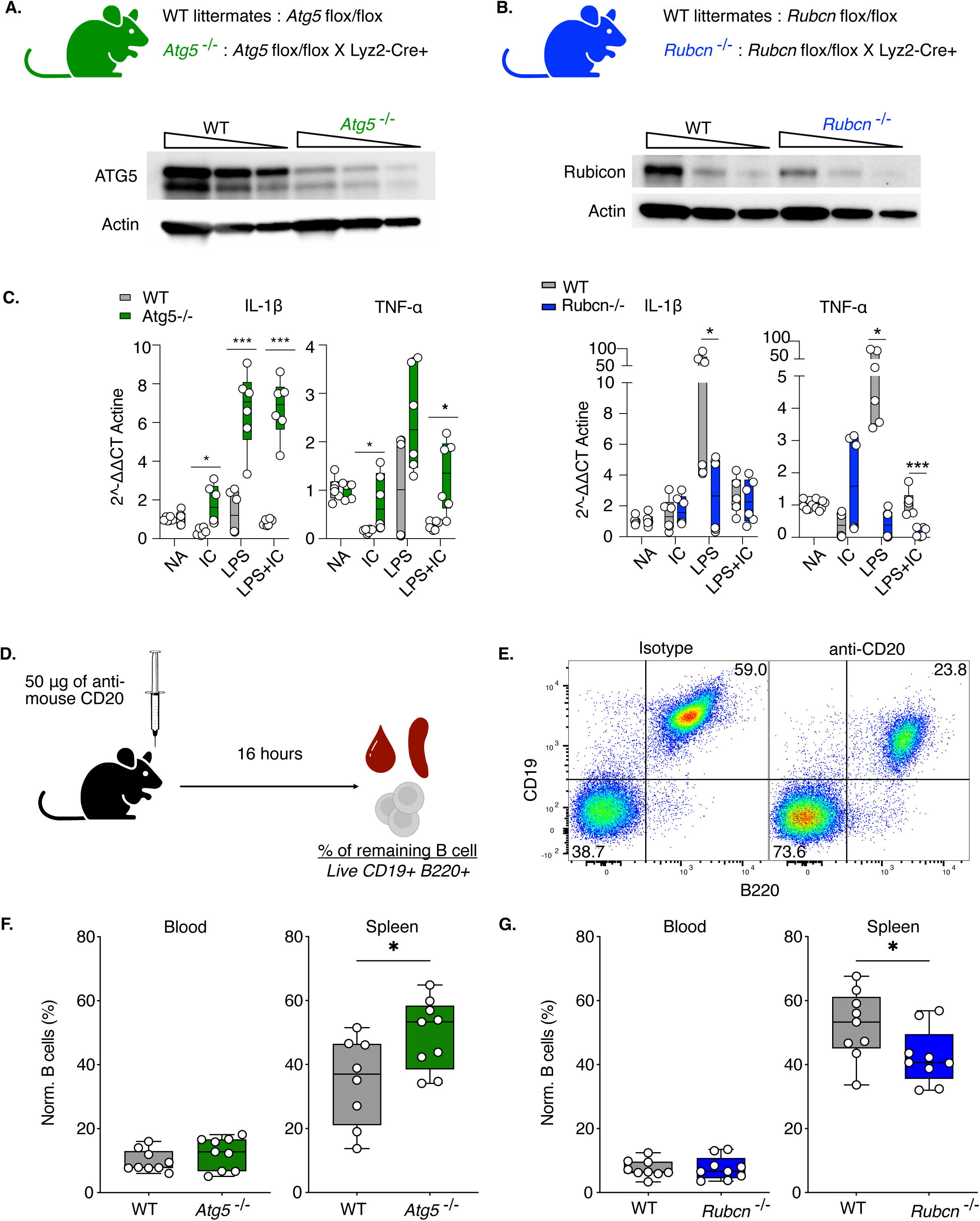
Autophagy is involved in FcyR function *in vitro* and *in vivo*. (**A**) Atg5 -/- mice were generated by crossing Atg5 lox/lox mice with Lyz2-Cre+ mice. Immunoblot analysis of Atg5 in Atg5 -/- and Atg5 +/+ wild-type BMDMs. (**B**) Rubcn -/- mice were generated by crossing Rubcn lox/lox mice with Lyz2-Cre+ mice. Immunoblot analysis of Rubicon in Rubcn -/- and Rubcn +/+ wild-type BMDMs. (**C**) Bone-marrow derived macrophages (BMDMs) were incubated without (NA) or with mouse IgG and anti-mouse IgG at 4°C. After removal of excess antibodies, the cells were incubated at 37°C in a normal or LPS-containing medium for 4.5h. Cells were then lysed, RNA extraction, RT and qPCR were performed according to kit instructions. The graphs show the result in 2^DDCt compare to actin. Experiments were performed 2 times. (**D**) Schematic representation of the B cell depletion model used to study FcyR function *in vivo*. (**E**) Gating strategy of splenic B cells 16h after isotype or anti-CD20 mAb injection. (**F-G**) Wild-type, Atg5 -/- or Rubcn -/- mice were injected i.v. with anti-CD20 mAb and the proportion of remaining B cells in the blood and spleen was determined 16h after injection by flow cytometry. The graphs show the percentage of B cells remaining in each group normalized to the percentage of B cells in the isotype-treated group. n=9 mice per each group.

Given the complexity of autophagy-driven pathophysiological mechanisms in arthritis, we addressed an alternative, better understood model to evaluate FcyR function, antibody-dependent cellular phagocytosis (ADCP). ADCP is a key mechanism underlying the efficacy of anti-tumor monoclonal antibody therapies that require target cell deletion (Beers *et al*, 2016). The major ADCP effector cells in mice belong to the myeloid lineage (Uchida *et al*, 2004). To evaluate ADCP *in vivo*, we injected an anti-mouse CD20 monoclonal antibody, which induces B cell depletion (Figure 3D-E) (Carter *et al*, 2017; Gillis *et al*, 2017). Under these conditions, peripheral blood B cell depletion, which is evoked mainly by Kupffer cells (Gong *et al*, 2005; Montalvao *et al*, 2013), was not affected by autophagy modulation (Figure 3F-G). In contrast, splenic B cell depletion was decreased in Atg5 -/- mice and increased in Rubcn -/- mice (Figure 3F-G), suggesting that autophagy is involved in ADCP mechanisms in splenic myeloid cells.

### The impact of autophagy modulation on FcyR trafficking, expression and signaling

Since autophagy is tightly linked to endocytic trafficking (Yamamoto *et al*, 2023), we explored whether altered endosomal trafficking of Fc receptors and ICs could underly FcyR-mediated ADCP in the spleen. To test this hypothesis, we used confocal microscopy to examine IC internalization and colocalization with EEA1, a marker of early endosomes, IRAP, a marker of storing signaling endosomes and LAMP1, a marker of lysosomes. In Atg5-/- macrophages, IC colocalization with EEA1 was not altered, but IC were retained for longer in IRAP endosomes (Supplementary Figure 2A-D) and were not targeted to LAMP1-positive vesicles (Figure 4A-B). In the presence of chloroquine, an inhibitor of lysosomal degradation, ICs could be detected in lysosomes at early time points, but their lysosomal localization in Atg5 -/- cells was less pronounced at later time points compared to wild-type cells (Figure 4C-D). In the absence of Rubicon, 15 minutes after internalization, IC colocalization with EEA1 was decreased, while the colocalization with IRAP was slightly increased (Supplementary Figure 3A-D). In contrast with Atg5 deficiency, Rubicon deficiency did not affect the IC targeting to lysosomes (Figure 4E-F).

**FIGURE 4:**
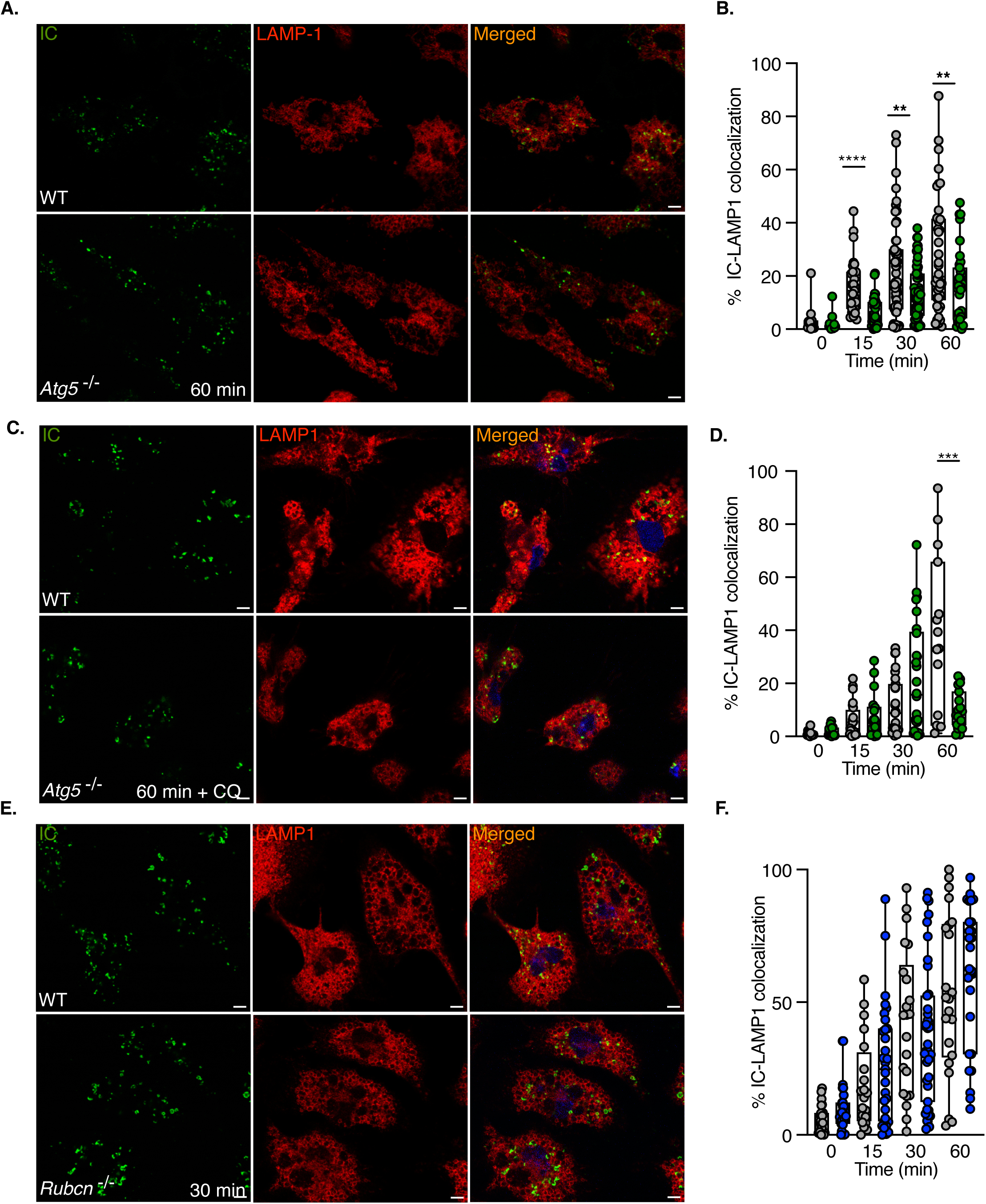
Autophagy modulation affects lysosomal targeting of immune complexes. (**A-B**) Atg5 -/- or wild-type BMDMs were incubated with mouse IgG and anti-mouse IgG conjugated to Alexa-Fluor 647 (green) at 4°C. After removal of excess antibody, cells were incubated at 37°C for 30 minutes, fixed, and stained with antibodies specific for LAMP-1 (red). (**C-D**) Atg5 -/- or wild-type BMDMs were incubated with mouse IgG and anti-mouse IgG conjugated to Alexa-Fluor 647 (green) at 4°C. After removal of excess antibodies, chloroquine was added and the cells were incubated at 37°C for 30 minutes, fixed and stained for LAMP-1 (red). (**E-F**) Rubcn -/- or wild-type BMDMs were incubated with mouse IgG and anti-mouse IgG conjugated to Alexa-Fluor647 (green) at 4°C. After removal of excess antibody, cells were incubated at 37°C for 30 minutes, fixed and stained for LAMP-1 (red). The figures show representative images from 2 independent experiments and the graphs show the percentage of ICs that colocalized with EEA1, IRAP or LAMP-1 within a cell. Each dot represents one cell. Scale bars = 5 μm.

We next evaluated whether changes in endosomal trafficking induced by the absence of Rubicon or Atg5 may impact FcyR expression in spleen phagocytes (CD64+/MertK+). While Rubicon deletion had no effect on FcyR expression at the plasma membrane, as assessed by flow cytometry, Atg5 deletion led to higher expression of FcyRIV, but not FcyRI or FcyRIII (Figure 5A). Increased expression of the high affinity FcyRIV would be expected to result in elevated ADCPas was demonstrated after anti-CD27 mAb stimulation in our prior studies (Turaj *et al*, 2017) but this was not observed here. These results demonstrate that the expression level of FcyRs cannot explain the effect of Atg5 or Rubicon deletion on ADCP function.

**FIGURE 5:**
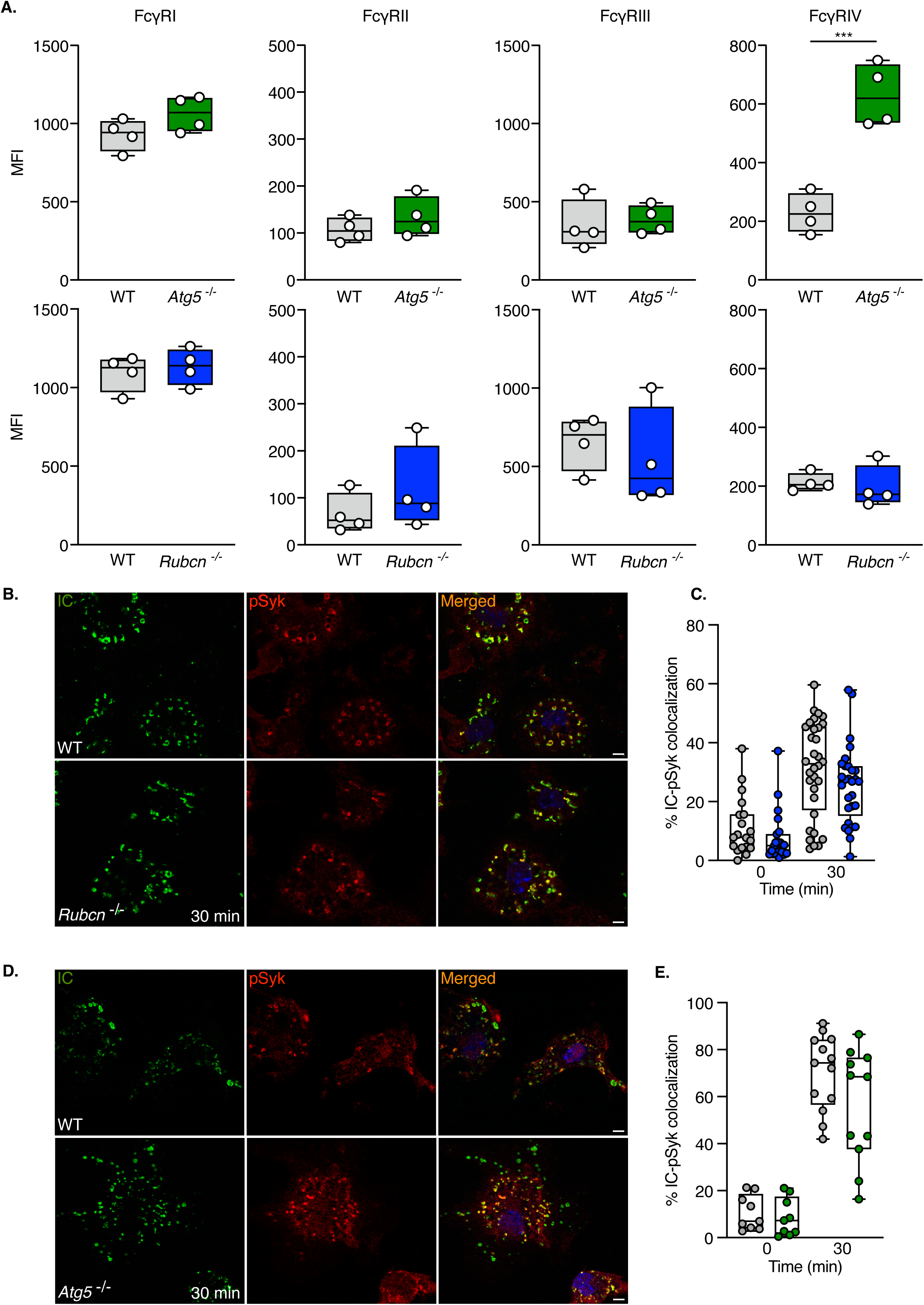
Autophagy modulation does not affect the ability of FcyRs to recruit Syk. (**A**) Mean fluorescence intensity (MFI) of each FcyR in MertK+ FcyRI+ splenic phagocytes from Atg5 -/-, Rubcn -/- or their respective wild-type littermates. n=4 mice for each group. (**B-E**) Atg5 -/-, Rubcn -/- or wild-type bone marrow derived macrophages (BMDMs) were incubated with mouse IgG and anti-mouse IgG conjugated to Alexa-Fluor647 (green) at 4°C. After removal of excess antibodies, cells were incubated at 37°C for 30 minutes, fixed and stained for pSyk (red). The pictures show representative images from three independent experiments and the graphs show the percentage of ICs that colocalized with pSyk within a cell. Each point represents one cell. Scale bars = 5 μm.

However, autophagy may affect the ability of the receptors to recruit the signaling machinery, including the key Syk protein kinase that associates with the receptor not only at the plasma membrane but also in IRAP storage endosomes (Benadda *et al*, 2023). Since the internalization of ICs into IRAP storage endosomes was slightly altered by deletion of both Rubicon and Atg5, we investigated whether the colocalization of ICs with the active form of Syk at the endosomal level is affected by modulation of autophagy. This was not the case, as both Rubicon- and Atg5-deficient cells showed the same level of recruitment of active Syk at the endosomal level as wild-type cells (Figure 5B-E). In conclusion, despite moderate changes in endosomal trafficking, modulation of autophagy partially affected the expression of FcyR but not their association with the active form of Syk, excluding the possibility that autophagy affects ADCP through reduced FcyR signaling.

### Autophagy modulation affects the monocyte transcriptomic landscape and reduces the number of monocytes

In an attempt to understand why changes in autophagy affect ADCP capacity, we performed a transcriptomic analysis of splenic phagocytes deficient for either Atg5 or Rubicon, isolated using MertK and FcyRI expression (Figure 6A). Neither Rubicon, nor Atg5 deletion affected the percentage of MertK+/ FcyRI+ cells in the spleen (Figure 6B). However, Atg5 deletion had a profound impact on gene expression, with 598 down-regulated and 334 up-regulated transcripts (Figure 6C and Supplementary Table 1). In contrast, Rubicon deletion had a modest effect on gene expression, with only 126 genes down-regulated and 34 genes up-regulated (Figure 6D and Supplementary Table 2). Among the few pathways significantly upregulated in the absence of Rubicon was the cellular response to interferons (IFN) (Supplementary Figure 4A-B), which may explain the more efficient ADCP in Rubicon-deficient cells, as both type I and type II IFN have been shown to increase the ability of macrophages to perform ADCP (Fan *et al*, 1991).

**FIGURE 6:**
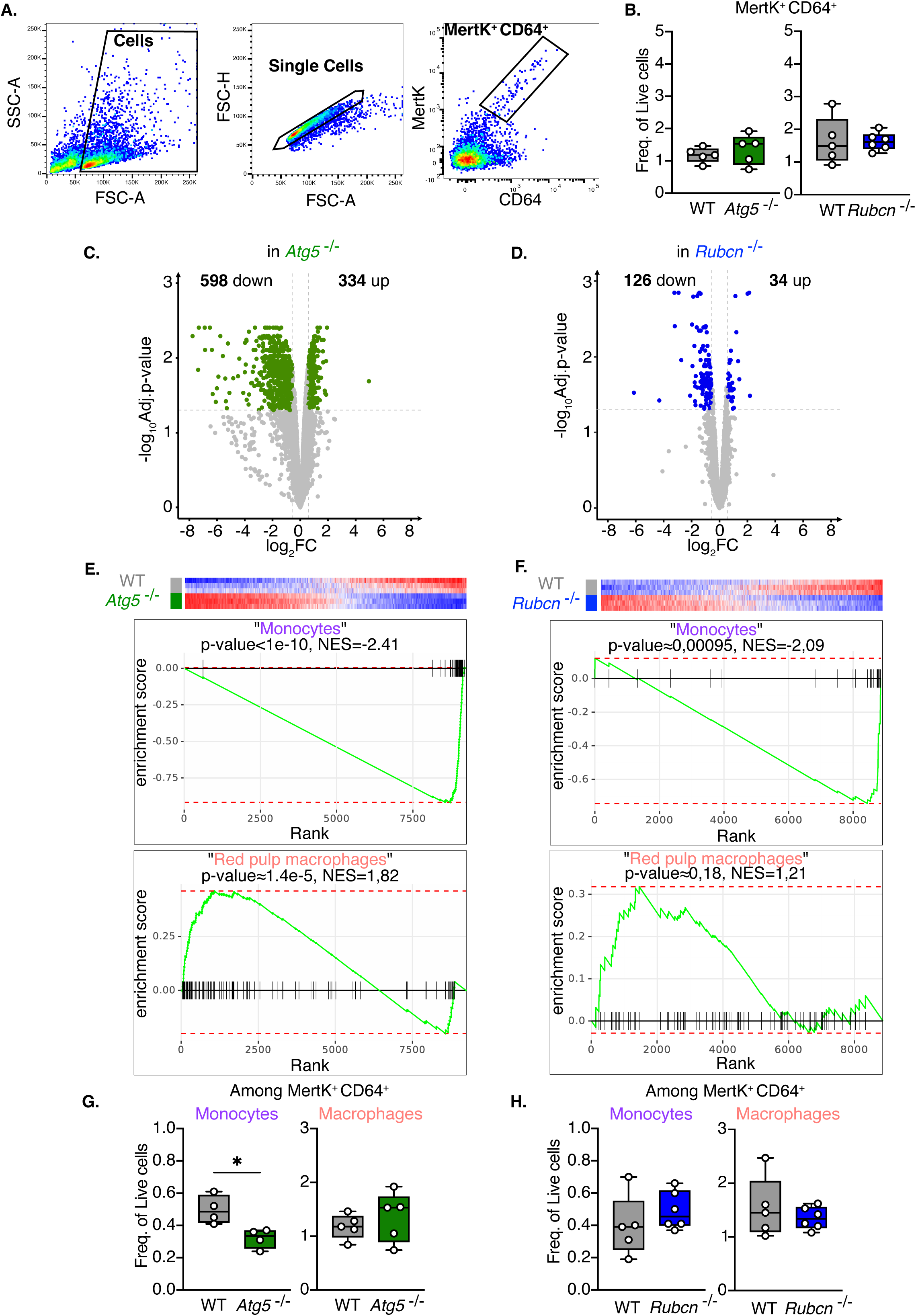
Atg5 deletion affects the monocyte signature in splenic phagocytes. (**A**) Gating strategy used to sort Mertk+ FcyRI+ splenic phagocytes for RNA sequencing analysis. **(B)** The graphs show the frequency of Mertk+ FcyRI+ splenic phagocytes among live cells. n=5 mice in each group. (**C-D**) Volcano plots showing genes up- and downregulated in Atg5 -/- or Rubcn -/- Mertk+ FcyRI+ spleen phagocytes compared to their wild-type counterpart. Genes are considered differentially expressed if they meet the following 2 criteria: Adjusted p-value <0.05 and log_2_FC>0.58 or <-0.58. (**E-F**) Gene set enrichment analysis of "monocytes" and "red pulp macrophages" on Mertk+ FcyRI+ splenic phagocytes from Atg5 -/-, Rubcn -/- or their wild-type counterparts. (**G-H**) The graphs show the frequency of monocytes and macrophages among Mertk+ FcyRI+ splenic phagocytes. n=5 mice in each group.

By analyzing the downregulated pathways in Atg5-deficient cells, we found that cytoplasmic protein synthesis was affected (Supplementary Figure 4C-D), suggesting a potential defect in cell development, fitness or function. Despite this indication, the percentage of MertK+/ FcyRI+ cells was identical in the absence of autophagy (Figure 6B), which may be explained by the presence of several cell populations in the MertK+/ FcyRI+ gate, the most numerous of which should be red pulp macrophages and blood-derived monocytes. To investigate whether monocyte or red pulp macrophage populations were differentially affected by autophagy changes, we defined signatures using previously published datasets (Heng *et al*, 2008), identifying the top 100 genes differentially expressed between monocytes and red pulp macrophages. Gene set enrichment analysis (GSEA) revealed a significant reduction in the monocyte signature and a significant enrichment in the red pulp macrophage signature in Atg5-deficient samples (Figure 6E and Supplementary Figure 5A), whereas the red pulp macrophage signature was not different in Rubicon-deficient samples compared to wild-type samples (Figure 6F and Supplementary Figure 5B). Of note, the GSEA was statistically significant for the reduction of monocyte signature in Rubicon-deficient samples, but this was based on a relatively small number of genes, 25, of which only 10 showed a significant adjusted p-value (Figure 6F and Supplementary Figure 5B). In summary, the results of the GSEA indicated a decrease in monocytes in Atg5-deficient samples and possibly also in Rubicon-deficient samples. To validate these GSEA results, we further refined the analysis of MertK+ FcyRI+ splenic phagocytic populations by excluding neutrophils, (identified as Ly6G++ cells), and gating CD11b-positive cells as monocytes and CD11b-negative cells as macrophages (Supplementary Figure 6). Using this gating strategy, it was apparent that in the absence of autophagy, the number of monocytes in the spleen was significantly reduced (Figure 6G), whereas Rubicon-deficient mice had no difference in their splenic monocyte numbers (Figure 6H).

Since the absence of autophagy affected the number of monocytes in the spleen (Figure 6G-H), we analyzed how autophagy modulates the transcriptomic landscape of macrophages and monocytes isolated from the spleens of Atg5-deficient mice. We sorted monocytes as Ly6G negative, CD11b positive and MHCII negative cells and macrophages as Ly6G negative, CD11b negative cells co-expressing F4/80 and FcyRI (Supplementary Figure 7A). Our initial analysis confirmed that our gating strategy did indeed isolate monocytes and macrophages, whose transcriptomes are very different (Figure 7A-B, Supplementary Figure 7B). Looking within each cell population, we found that the absence of autophagy strongly affected the transcriptome of monocytes with a less profound impact on macrophages (Figure 7C-D and Supplementary Tables 3-5). The fact that autophagy has a limited effect on splenic macrophages in our study (Figure 7D) may be explained by the promoter used to delete Atg5. Lyz2 is very well expressed in monocytes, whereas its expression in splenic resident macrophages is very low (Supplementary Figure 7C). Finally, the analysis of the pathways affected by Atg5 deletion in myeloid cells showed that these cells downregulate genes involved in DNA metabolism and cell cycle regulation and upregulate genes involved in the inflammatory response (Figure 7E-F) and alternative protein degradation pathways upon cellular stress, such as endoplasmic reticulum-associated protein degradation, ERAD (Supplementary Figure 7D).

**FIGURE 7:**
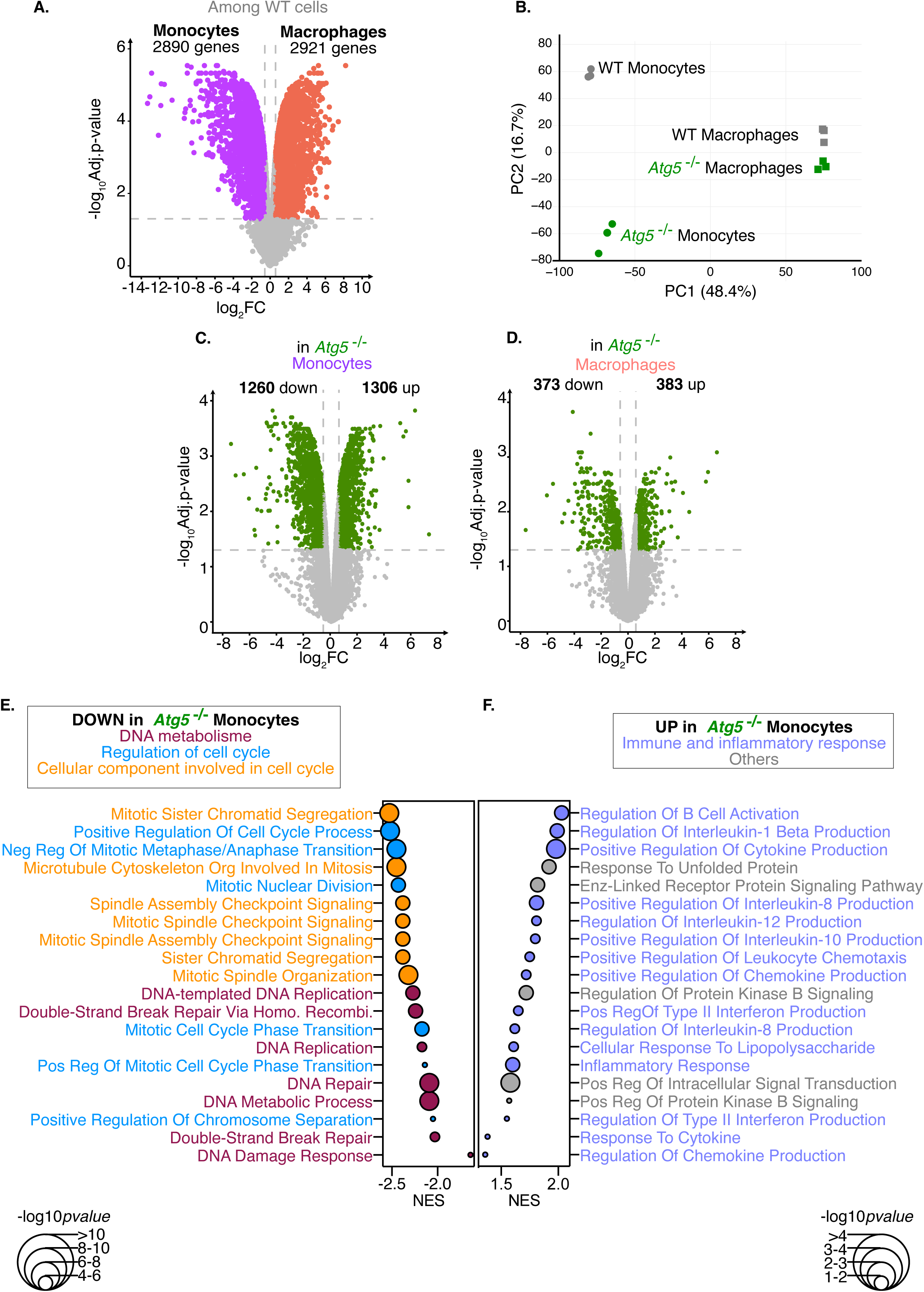
Autophagy-deficient monocytes show cell cycle abnormalities and an increase in the inflammatory response. (**A**) Volcano plot showing genes differentially expressed between wild-type monocytes and macrophages. Genes are considered differentially expressed if they meet the following 2 criteria: Adjusted p-value <0.05 and log_2_FC>0.58 or <-0.58. **(B)** Principal component analysis (PCA) of splenic monocytes or macrophages obtained from 3 different wild-type or Atg5 -/- mice. (**C-D**) Volcano plots represent genes up- and downregulated in Atg5-/- monocytes or macrophages compared to their wild-type counterpart. Genes are considered differentially expressed if they meet the following 2 criteria: Adjusted p-value <0.05 and log2FC>0.58 or <-0.58. (**E**) The graph shows the Normalized Enrichment Score (NES) and its -log_10_p value of the top 20^th^ down-regulated pathway in Atg5 -/- monocytes. Dark red pathways are associated with DNA metabolism, blue pathways are associated with cell cycle regulation, and orange pathways are associated with cellular components involved in the cell cycle. (**F**) The graph shows the Normalized Enrichment Score (NES) and its -log_10_p value of the top 20th upregulated pathway in Atg5 -/- monocytes. Purple pathways are associated with immune and inflammatory response.

Considering that cell proliferation pathways were affected by Atg5 deletion (Figure 7E), we specifically looked at cell proliferation markers, such as the nuclear antigen for cell proliferation (PCNA) and the proliferation marker Ki67. The expression of these two genes was significantly decreased in the absence of autophagy (Supplementary Figure 7E). Furthermore, among the transcription factors involved in monocyte differentiation, we observed a significant decrease in the transcription factor *Cebpe* (Supplementary Figure 7E), which is involved in the terminal differentiation of monocytes (Huber *et al*, 2014). All these transcriptomic results, as well as the decrease in monocytes observed by flow cytometry (Figure 6G), show that autophagy is important for the differentiation and survival of monocytes, as previously reported for other cell types (Das *et al*, 2012).

In conclusion, our study shows that autophagy is induced by FcyRI activation, but at the same time it is required for the differentiation of monocytes, which, thanks to their FcyRs, participate in ADCP, an essential mechanism for therapeutically important monoclonal antibodies.

## Discussion

In this work, we show that similar to other innate immune receptors, such as TLRs and NOD-like receptors, activation of FcyRs induces canonical autophagy. Interestingly, the induction of autophagy downstream of innate immune receptor activation appears to be receptor specific. For example, TLR4 induces autophagy via the TRIF-p38 axis and not via MyD88 (Xu *et al*, 2007), whereas TLR7 induces autophagy via MyD88 (Delgado *et al*, 2008), and TLR2 induced autophagy requires ERK activity (Anand *et al*, 2011). Our data show that for FcyRs, their cross-linking leads to the recruitment of the phosphatase Pp2a, one of the two phosphatases capable of dephosphorylating ULK1 at the Ser637 residue and inducing autophagy (Wong *et al*, 2015). Recruitment and activation of Pp2a by signaling cascades downstream of the BCR, another ITAM-coupled immune receptor that can trigger autophagy, has been reported (Inui *et al*, 1998). In the case of the BCR, the phosphatase interacts directly with immunoglobulin binding protein 1 (Igbp1). Whether Igbp1 is also involved in phosphatase recruitment and activation downstream of FcyRs remains to be investigated.

Autophagy activation had multiple effects on myeloid cell function. First, we showed that autophagy limits the amplitude of the FcyR-induced inflammatory response, as demonstrated by an increased production of IL-1β and TNF-α in Atg5-deficient macrophages. This anti-inflammatory role of autophagy is consistent with previously demonstrated anti-inflammatory roles of autophagy, such as the inhibition of retinoic acid-inducible gene I (RIG-I) and STING signaling or prevention of inflammasome activation by removal of damaged mitochondria (Deretic *et al*, 2013). Whether the role of autophagy in limiting the FcyR-induced pro-inflammatory response is relevant *in vivo* remains an open question. We sought to analyze a model of FcyR-induced disease by inducing arthritis with a cocktail of anti-collagen antibodies and LPS. This model was unaffected by modulation of autophagy (through loss of Atg5 or Rubicon) in Lyz2-expressing myeloid cells. The most plausible explanation for these results is the high expression of Lyz2 in osteoclasts, which are thus autophagy deficient in Atg5^loxlox^ x Lyz2-Cre mice. Since osteoclast differentiation is impaired in the absence of autophagy, they may no longer have a negative impact on bone erosion and disease development (Lin *et al*, 2013). As in the case of osteoclasts (Lin *et al*, 2013), our results demonstrated that autophagy deletion in monocytes affected monocyte numbers and fitness.

The ADCP defect observed in mice lacking autophagy in myeloid cells may be either a consequence of the reduced number of monocytes. However, it is also possible that in addition to cell differentiation and survival, autophagy controls the production of ADCP effector molecules. Looking at the effectors of macrophage cytotoxicity, we found in our transcriptome data that the only factor that was significantly reduced in the absence of autophagy was the antimicrobial peptide CRAMP (Supplementary Figure 7F), whose involvement in antibody dependent target cell killing has been reported (Bruns *et al*, 2015). Our *in vivo* ADCP experiments demonstrated that the anti-CD20 mAb-mediated depletion of peripheral blood B cells was unaffected in autophagy deficient mice, in contrast to the defect in the depletion of splenic B cells in the same Atg5-/- mice. This finding may be explained by the involvement of different myeloid cells in B cell depletion in the two organs. B cell depletion in the blood is performed by liver macrophages, of which Kupffer cells are the major population (Montalvao *et al*, 2013). Our results suggest that ADCP mechanisms may differ between splenic myeloid cells and liver macrophages, which is perhaps unsurprising given the strong tissue imprinting of macrophages (Guilliams & Svedberg, 2021). Kupffer cells are derived from the embryonic Yolk sac and in the adult they principally clear cell debris, in an immunologically silent manner, being dedicated to immune tolerance (Mass *et al*, 2023). Splenic myeloid cells, by contrast, include blood-derived monocytes as well as Yolk sac-derived resident macrophages, among which the most abundant are red pulp macrophages, responsible for the immunologically silent clearance of blood cells (Zhao *et al*, 2024). The Cre expression driven by the Lyz2 promoter used in our study should strongly modulate autophagy in blood-derived monocytes (Supplementary Figure 7C) and liver Kupffer cells (Sakai *et al*, 2019) and to a lesser extent red pulp macrophages (Supplementary Figure 7C). Thus, both blood-derived spleen monocytes and Kupffer cells should be autophagy-deleted, but the autophagy deletion did not affect the ability of Kupffer cell to perform ADCP. This may be due to alternative mechanisms of ADCP in Kupffer cells, which needs further investigation.

Finally, our study opens tempting perspectives for the modulation of autophagy in the context of therapeutic use of monoclonal antibodies and also in cellular immunotherapy. Thus, we can hypothesize that the use of monoclonal antibodies against tumor antigens could be combined with rapamycin or other mTOR inhibitors to enhance autophagy induction. In support of this hypothesis, mTOR inhibitors have already been tested in combination with anti-tumor monoclonal antibody therapy and were shown to improve treatment efficacy (Xu *et al*, 2014; Miller *et al*, 2009). However, the mechanism of this improvement has not been elucidated, and in light of our results, an increase in ADCP efficacy could be involved. Along the same line, PP2A-activating drugs have been shown to be efficient in anti-tumor therapy, but the majority of these studies did not investigate the effect of PP2A-activating drugs on myeloid cells, they mainly focused on the direct role of PP2A in suppressing tumor cell growth (Neviani *et al*, 2013). Nevertheless, the potential use of PP2A-activating drugs in inflammatory diseases has been proposed, highlighting the role of the phosphatase as an anti-inflammatory factor, but without linking this effect to autophagy induction (Clark & Ohlmeyer, 2019). Last but not least, it is important to consider our results in the view of the development of chimeric antigen receptor (CAR)-carrying macrophages, a cellular immunotherapy, that, unlike CAR T cells, appears to have the potential to infiltrate solid tumors (Abdin *et al*, 2024). CAR macrophages could, at least in theory, take full advantage of increased autophagy, either through genetic inactivation of Rubicon, which in our study increased the efficacy of ADCP, or through combinations with mTOR inhibitors such as rapamycin, which induce activation of autophagy.

## Methods and protocols

### Cell culture

BMDMs were generated *in vitro* by culturing bone marrow precursor cells from large bones for 7 days in complete medium (IMDM supplemented with 10% FCS, 2 mM glutamine, 100 U/mL penicillin, 100 μg/mL streptomycin, 50 μM β-mercaptoethanol supplemented with 20% of L929-conditioned media containing M-CSF). RAW macrophages were cultured in complete medium (DMEM high glucose supplemented with 10% FCS, 2 mM glutamine, 100 U/mL penicillin, 100 μg/mL streptomycin). DC2.4-hCD32a-GFP-APEX2 were cultured in complete medium (IMDM supplemented with 10% FCS, 2 mM glutamine, 100 U/mL penicillin, 100 μg/mL streptomycin, 50 μM β-mercaptoethanol). All the experiments were performed in the presence of 10% FCS.

### Mice

Rubcn ^lox/lox^ mice were a generous gift from Dr. T Takehara (Tanaka *et al*, 2016)and Atg5 ^lox/lox^ x LysM-Cre from Dr. S. Lotersztajn (Lodder *et al*, 2015). All mice were genotyped to verify presence or absence of LysM-cre, Atg5^lox/lox^ and Rubcn^lox/lox^ using specific primers. Mice were bred in a conventional facility, with ambient temperature between 20 and 24 °C, humidity between 40 and 60%, and a continuous light–dark cycle. Both male and female mice between 8 and 14 weeks of age were used in the experiments. All animal experiments were approved by the Comité d’éthique pour l’expérimentation animale Paris-Nord/N° 121 (APAFIS#31323-2021042612121238), additionally approved by the CRI U1149 Ethical Committee and by the French OGM Committee (DUO n° 5643).

### Receptor aggregation for confocal microscopy, proximity ligation assay and immunoblot

For immunoblot, 1 million BMDMs were plated on a 6-well plate and incubated overnight. For confocal microscopy and proximity ligation assay, 50.000 RAW cells and BMDMs were seeded on fibronectin-coated slides for 16 h. To cross-link FcγRs, cells were incubated with mouse IgGs (10 μg/mL) for 15 min on ice. Cells were washed twice in PBS and receptor cross-linking was completed by adding a donkey anti-mouse Alexa Fluor 647 (10 μg/mL). Cells were washed twice in PBS and then incubated to 37°C for the time points indicated in the assays. All the incubations at 37°C were performed in complete medium, with 10% FCS, with chloroquine (50 uM), or with 2.5 µM Pp2a inhibitor (LB100), as required. For experiments with LB100, cells were incubated with the inhibitor for 2 hours prior to FcyR cross-linking.

### Immunofluorescence microscopy

Cells were fixed with 2% PFA for 15 minutes at 37°C. Cells were then permeabilized with 0.2% saponin in PBS containing 0.2% BSA. Staining was performed in the same buffer by incubating 40 µL of diluted primary antibody in a humidity chamber for 40 minutes at room temperature (RT), followed by two washes in the staining buffer. The slides were then incubated in 40 µL of 1:100 diluted secondary antibody conjugated either with AF488 or AF594. After two washes in the staining buffer and an additional wash in PBS, cells were postfixed in 4% formaldehyde for 10 minutes at RT, washed in PBS and quenched in NH_4_Cl (5mM). Slides were washed again and incubated with DAPI in PBS for 5 minutes before mounting in a Fluoromont-G drop. Images were acquired on a Leica SP8 confocal microscope, with a 63x oil immersion objective. Image treatment and analysis were performed using ImageJ software. Marker colocalizations were evaluated using only unsaturated images and the ImageJ software. A manual threshold was set for each channel before image analysis. Individual cells were selected using the freehand selection tool and considered as a region of interest (ROI) in ImageJ. For colocalization studies, all images were first converted to a binary image (black pixel intensity = 0; white pixel intensity = 1). The binary images for independent channels were multiplied to create a mask that included the pixels present in both channels. The areas of the pixels for each color and the mask pixels were calculated using ImageJ’s measure stack plugin. The percentage of pixels of one color that colocalized with pixels of another color was calculated as the ratio of the sum of the area of the pixels in the mask divided by the sum of the area of the pixels of the first color. Statistical analysis was performed with GraphPad Prism software using unpaired t-tests. For p62 dots, p62 dots were quantified using a particle size MACRO based on ImageJ. In the colocalization and p62 dot graphs, each dot represents one cell.

### Proximity ligation assay

Cells were fixed with 2% PFA for 15 minutes at 37°C. The cells were then permeabilized with 0.5% Triton 10x in PBS. Staining was performed in PBS-0.2% BSA by incubating cells with 40 µL of diluted primary antibody in a wet chamber for 2 hours at RT, followed by two washes in staining buffer. Duolink™ (OLINK Bioscience) was performed according to the manufacturer’s instructions. After washing, PLA probes were added, followed by hybridization, ligation and amplification for 100 minutes at 37°C. Protein interactions were visualized after incubation with red detection solution. Before mounting in a Fluoromont-G drop, the slides were washed again and incubated with DAPI in PBS for 5 minutes. Images were captured on a Leica SP8 confocal microscope with a 63x oil immersion objective. Image processing and analysis were performed using ImageJ software, and quantification of the dots was performed using a particle size MACRO based on ImageJ.

### Immunoblotting

Cell pellets were resuspended in 1x Laemmli buffer. Samples were boiled at 95°C for 10 minutes, then the lysates were cleared by centrifugation at 15000g for 10 minutes. Proteins were separated by SDS-PAGE on Criterion 4–15% acrylamide gels (Bio-Rad) in Tris-Glycine-SDS buffer and transferred on PVDF or nitrocellulose membranes (Bio-Rad) using a Trans-Blot Turbo Transfer System from Bio-Rad. Membranes were blocked at least 1 hour in 5% BSA-TBST and incubated with each primary antibody overnight at 4°C, washed extensively in TBST buffer, and incubated 1 hour with secondary HRP-conjugated antibody at RT. After extensive washing, the membranes were incubated with Clarity Western ECL Substrate (Bio-Rad) for 5 min. The chemiluminescence signal was acquired using a ChemiDoc Imaging System and quantified using ImageLab software (Bio-Rad).

### In situ biotinylation

#### Cloning and expression of FcyRIIA-GFP-APEX2

The DNA fragment of CD32A-GFP was amplified from a donor plasmid by touch-down PCR and was cloned by recombination (In-Fusion HD Cloning Kit; 121416; Takara Bio USA) into the pHR vector already containing the APEX2 sequence, using the following primers:

Fw: 5’-gagaattctcaCgCgTgccaccATGGAGaCCCAAATGTCTCA-3’

Rev: 5’-catggacgagctgtacaagCgAT*cgcaccggtatacaa*-3’

The primers contain in their 5’ end a sequence that allows recombination to the recipient plasmid and they were designed under the guidelines of the manufacturer.

Briefly, the pHR vectror was linearized using AsiSI and MluI (R0630S; R3198S; New England Biolabs) for 2h at 37°C. The linearized vector and the PCR fragment were purified by gel extraction using a gel and PCR clean-up kit (740609.250; Macherey-Nagel). The recombination reaction was performed using 50ng vector DNA with a 2:1 insert:vector ratio under manufacturer’s guidelines. Chemically competent Stbl3 bacteria were transformed with 2.5μl recombination reaction and single colonies were isolated for DNA amplification. The recombination was verified by diagnostic digestion with EcoRI (R3101L; New England Biolabs) and the sequence was analyzed by Sanger sequencing (Eurofins). The lentiviral production and transduction of DC2.4 cells was performed as previously described (Benadda *et al*, 2023).

DC2.4-FcyRIIA-GFP-APEX2 were divided into 3 conditions, with respect to receiving Biotin, H_2_O_2_ and immune complexes (IC). Cells (50 million per condition) were incubated in 10 mL of medium containing biotin-phenol (500 µM) for 30 minutes at 37°C. Cells were then centrifuged at 500 g for 5 minutes at 4°C and washed once in ice-cold PBS. Receptor aggregation was then performed by adding 1mL of PBS containing 10 µg/mL of F(ab)’_2_ anti-FcyRIIA, followed by 15 minutes of incubation on ice, one wash with cold PBS, and then another 15 minutes of incubation on ice with 1 mL of PBS containing 10 µg/mL of F(ab)’_2_ anti-mouse F(ab)’_2_. The cells were washed once more, resuspended in Biotin-phenol containing medium, and incubated at 37°C for 30 minutes. Then, 10 mL of medium containing 2 mM H_2_O_2_ was added and the cells were incubated for 30 seconds before adding 10 mL of cold quenching buffer (10 mM sodium ascorbate, 10 mM sodium azide, 1 mM CaCl2, 1 mM Trolox in PBS). Cells were centrifuged and washed 3 times with 10 mL of quenching buffer, separated by centrifugation at 600 g for 3 minutes. Then, 10 mL of quenching buffer were added and the samples were incubated on ice for 20 minutes. After a final centrifugation step, the pellets were lysed in 250 µL of RIPA buffer (Tris-HCl 50 mM pH 7,5, 0,5% deoxycholate, 1% NP40, 10 mM sodium ascorbate, 0,10% SDS, 10 mM sodium azide, 1 mM DTT, 1 mM NaCl, 1 mM Trolox, 2,5 mM MgCl_2_ in H_2_O) containing DNAse I and 1mM protease inhibitors, for 20 minutes on ice with gentle shaking. The lysates were then centrifuged at 16000 g for 10 minutes, before being applied to a 2 mL Zeba Desalting 7 MWCO desalting column to remove any residual biotin-phenol, according to the Zeba Desalting kit instructions. 10 µL of each lysate were kept as “total cell lysate” for further blot analysis. The remaining cell lysate was incubated with 150 µL of magnetic streptavidin beads, which were previously washed with RIPA buffer. Samples were placed on a wheel for 2h 30 minutes at 4°C, before being washed 3 times in 1 mL of RIPA buffer using a magnetic rack. Finally, the beads were frozen for further analysis by immunoblot.

#### RT-qPCR

1 million of BMDM were plated on 6 wells plate overnight. To cross-link FcγRs, cells were incubated with mouse IgGs (10 μg/mL) for 15 min on ice. Cells were washed twice in PBS and receptor cross-linking was completed by adding a goat anti-mouse Alexa Fluor 647 (10 μg/mL). Cells were washed twice in PBS and then incubated to 37°C for 4 hours at 37°C, in complete medium, with 10% FCS, with or without LPS (5 ng/mL). Cells were washed and RNA extraction was performed following kit instruction (Macherey-Nagel, RNA Extraction). RT was performed with iScript cDNA synthesis kit (Bio-Rad) and qPCR was performed with OneGreen Fast qPCR mix (Ozyme).

#### Flow cytometry

Blood was collected in a 1.5 mL tube containing 20 µL of heparin. 3 cycles of red blood cell lysis as follows: first 1 mL of ACK, 10 minutes at 4°C, with second and third washed with 500 µL of ACK, 5 minutes, were performed on 50 µL of blood. These cycles were separated by addition of PBS and centrifugation for 5 min at 450 g, 4°C. After the last centrifugation, the cells were resuspended in FACS buffer (PBS, 2% FCS 2 mM EDTA) and distributed in a 96-well plate for staining.

Spleens were collected in a 6-well plate containing cold PBS. For B cell depletion analysis, spleens were smashed on a 70 µm filter and rinsed in PBS. For myeloid cell analysis, spleens were digested by injection of 500 µL of PBS containing 100 ug/mL DNAse and 200 ug/mL collagenase. The spleens were then cut into small pieces and incubated for 15 minutes at 37 °C under agitation. Mechanical digestions were performed by passing small pieces through an 18G needle, and samples were incubated for an additional 10 minutes at 37°C under agitation. Digestions were stopped by adding cold PBS and passing samples through a 70 µm filter, rinsed in PBS. Erythrocyte lysis was performed by adding 2 mL ACK for 2 minutes at RT after centrifugation at 500 g for 5 minutes. The reaction was stopped by adding cold FACS buffer and after a final step of 5 minutes centrifugation, the cells were resuspended in 1 mL FACS buffer and 1:10 of each spleen was added to a 96-well plate for staining. Data were acquired on a BD LSR Fortessa X20 using Diva software and analyzed using FlowJo software.

#### Cell sorting

Spleen samples were prepared the same way as described above. Cell sorting was performed on a BD Melody. 10,000 cells per sample were collected directly into 150 μl of TCL lysing buffer (Qiagen) containing 1% β-mercaptoethanol and immediately stored at - 80°C until RNA extraction. RNA was extracted using the Single Cell RNA purification kit (Norgen, Cat#51800) according to the manufacturer’s instructions. After extraction, total RNA was analyzed on an Agilent 2100 BioAnalyzer System. RNA quality was estimated using the RNA Integrity Number (RIN). RNA sequencing libraries were prepared using the SMARTer Stranded Total RNA-Seq Kit v2 - Pico Input Mammalian (Clontech/Takara). The input amount of total RNA varied between 1 and 10 ng per sample. This protocol includes a first step of RNA fragmentation, using a proprietary fragmentation mix at 94°C. The incubation time was set up for each sample, based on the RNA quality, and according to the manufacturer’s recommendations. After fragmentation, indexed cDNA synthesis was performed. The ribodepletion step was then performed, using probes specific for mammalian rRNA. Finally, PCR amplification was performed to amplify the indexed cDNA libraries, with a number of cycles set according to the amount of tRNA input. Library quantification and quality assessment were performed by Qubit fluorometric assay (Invitrogen) with dsDNA HS (High Sensitivity) Assay Kit and LabChip GX Touch using a High Sensitivity DNA Chip (Perkin Elmer). Libraries were then equimolarly pooled and quantified by qPCR using the KAPA library quantification kit (Roche). Sequencing was performed on the NovaSeq 6000 (Illumina), targeting between 15 and 20 M reads per sample and using paired-end 2 x 100 bp.

#### RNAseq analysis

Analysis of the raw data was performed using RStudio and subsequent analysis and visualization were performed using the Phantasus online tool doi:10.18129/B9.bioc.phantasus, https://ctlab.itmo.ru/phantasus). Differentially expressed genes were identified using a threshold of log2 fold-change (higher than 1.5 or lower than −1.5) and adjusted to a p value less than 0.05. Pathway analysis, Gene Set Enrichment Analysis (GSEA) and Normalized Enrichement Score (NES) were calculated based on the Gene Ontology Biological Process 2023 and Bioplanet 2019 databases.

#### CAIA model

Mice were injected intraperitoneally with 5 mg of a mix of 5 clone monoclonal anti-mouse type II collagen cocktail (Chondrex, #53100). 3 days later, mice were injected with 50 µg LPS. Joint thickness and arthritis score were measured daily. Mice were sacrificed on day 10.

#### B cell depletion

Mice were injected intravenously with 50 µg of anti-mouse CD20 (clone 5D2, Genentech). After 16 hours, blood and spleen were collected and processed as described above. The percentage of residual B cells was determined as the percentage of CD19- and B220-positive cells among live CD45+ cells. The normalized percentage of B cells corresponds to the percentage of B cells remaining after treatment compared to PBS-injected mice (100%).

## Supplementary Figure Legends

**Supplementary Figure 1:**
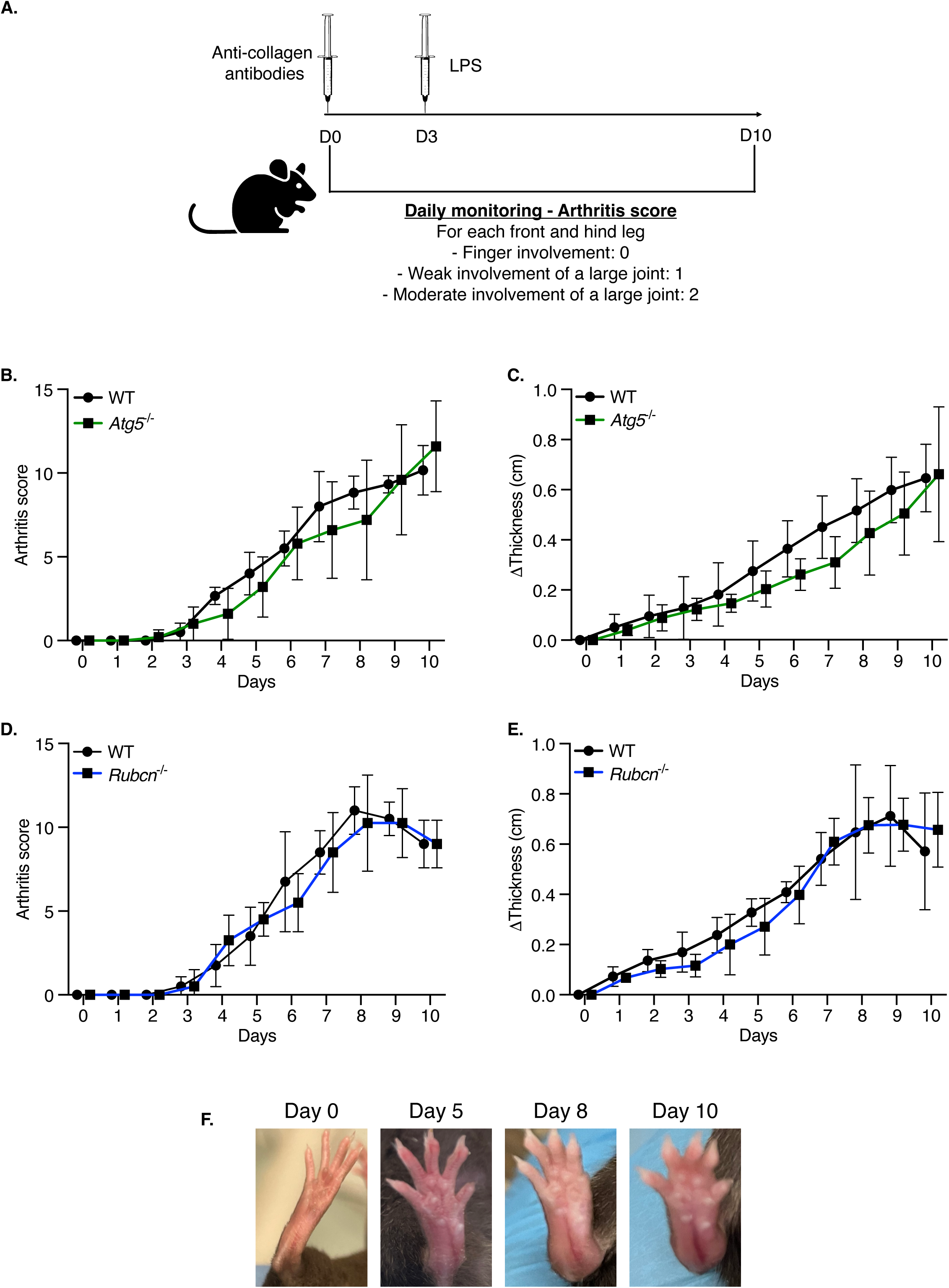
Autophagy modulation does not affect the pathophysiology of arthritis in a collagen antibody induced arthritis model. **(A) Schematic representation of the collagen antibody induced arthritis (CAIA) model** (**B-E**) The graphs show the arthritis score or mean delta joint thickness of Atg5 -/-, Rubcn -/- or their wild-type counterparts in CAIA arthritis model. n=5 mice per group. (**F**) The images show the hind paw and its development during the course of collagen antibody induced arthritis.

**Supplementary Figure 2:**
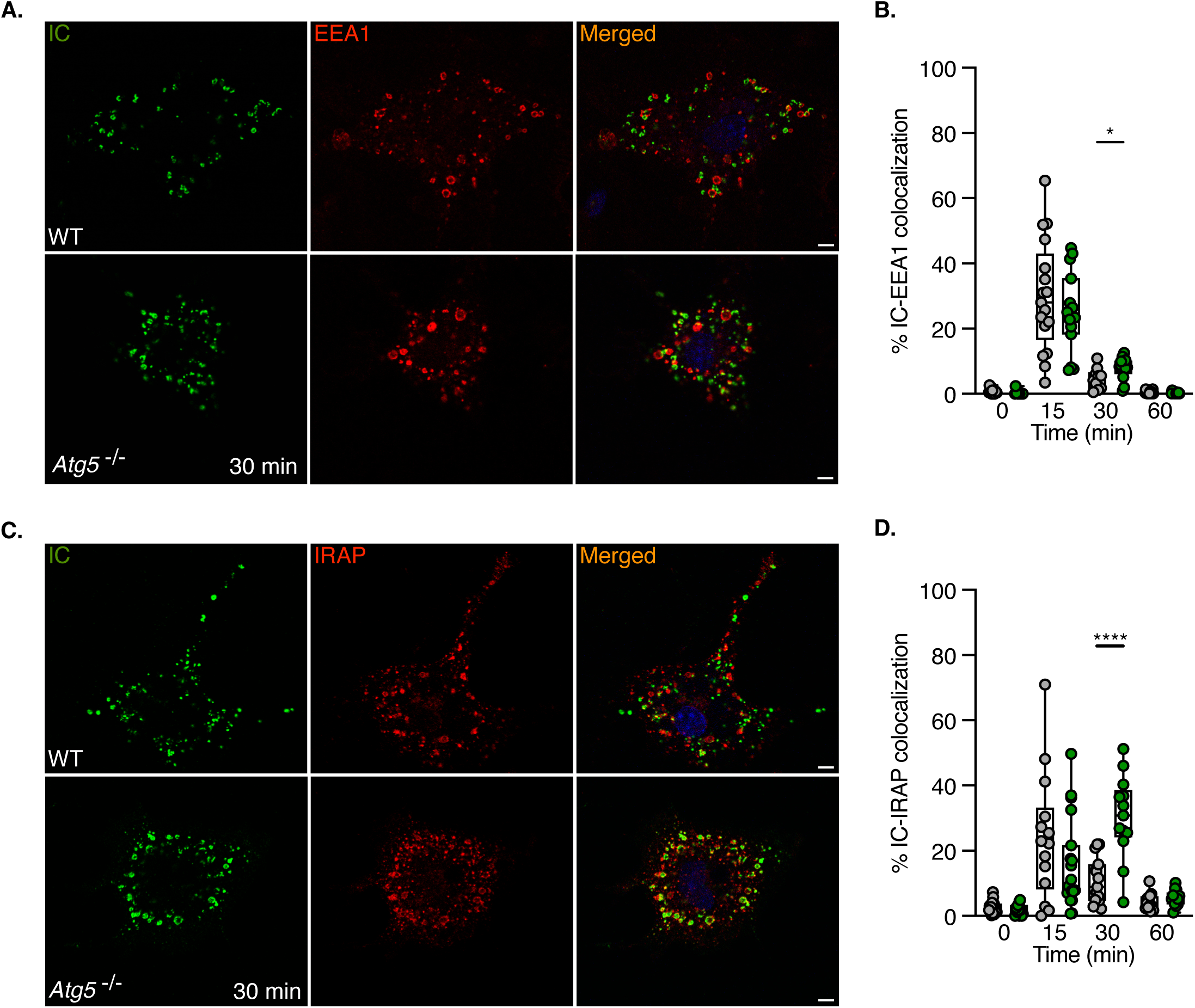
Atg5 deletion retains ICs in storage endosomes. (**A-D**) Atg5 -/- or wild-type BMDMs were incubated with mouse IgG and anti-mouse IgG conjugated to Alexa-Fluor 647 (green) at 4°C. After removal of excess antibody, cells were incubated at 37°C for 30 minutes, fixed, and stained with antibodies specific for EEA1 or IRAP (red). The figures show representative images from 2 independent experiments and the graphs show the percentage of ICs that colocalized with EEA1, IRAP or LAMP-1 within a cell. Each dot represents one cell. Scale bars = 5 μm.

**Supplementary Figure 3:**
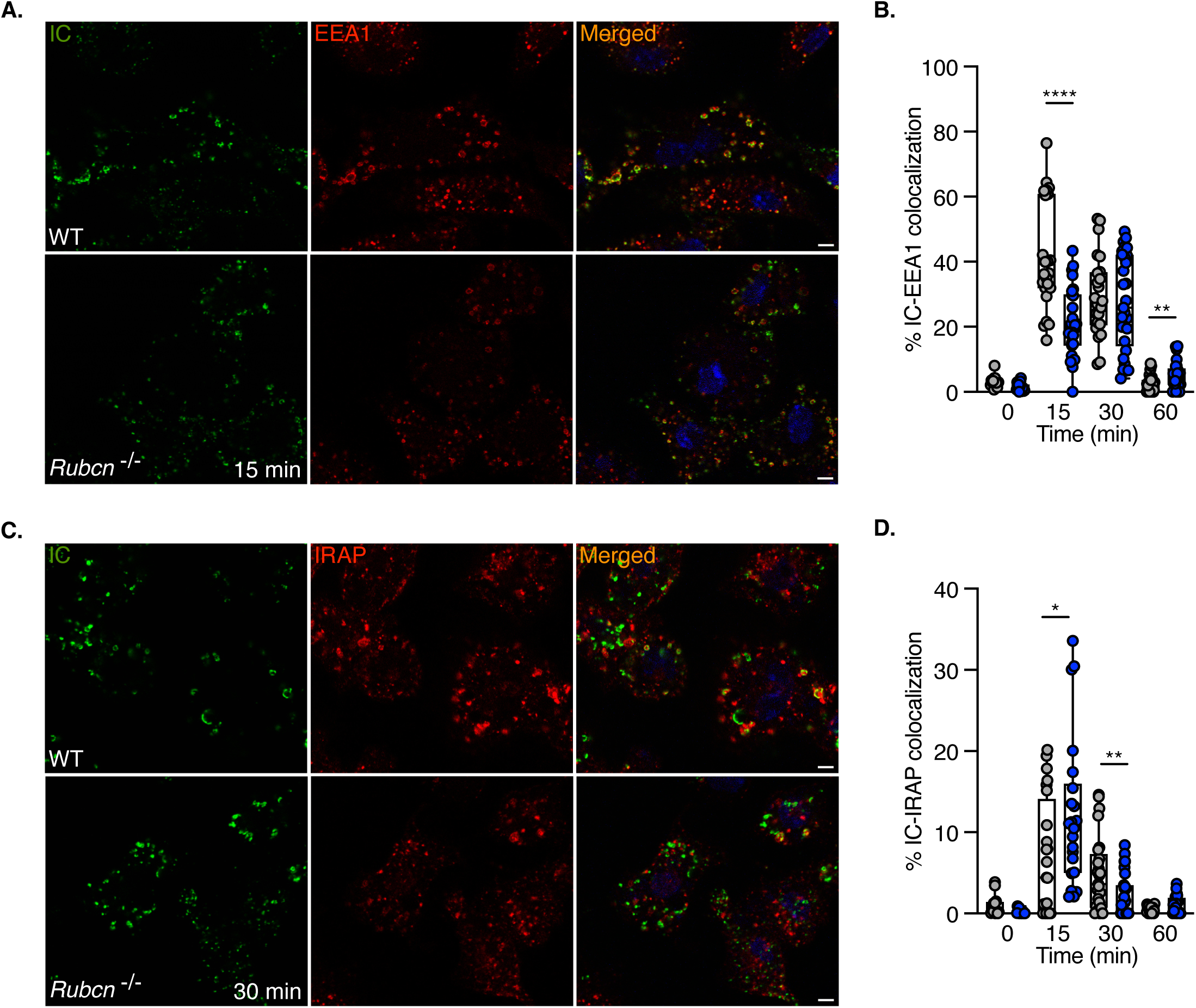
Rubicon deletion accelerates endosomal trafficking of ICs. (**A-F**) Rubcn -/- or wild-type BMDMs were incubated with mouse IgG and anti-mouse IgG conjugated to Alexa-Fluor647 (green) at 4°C. After removal of excess antibody, cells were incubated at 37°C for 30 minutes, fixed and stained for EEA1 or IRAP (red). The pictures show representative images from 2 independent experiments and the graphs show the percentage of ICs that colocalized with EEA1, IRAP or LAMP-1 within a cell. Each dot represents one cell. Scale bars = 5 μm.

**Supplementary Figure 4:**
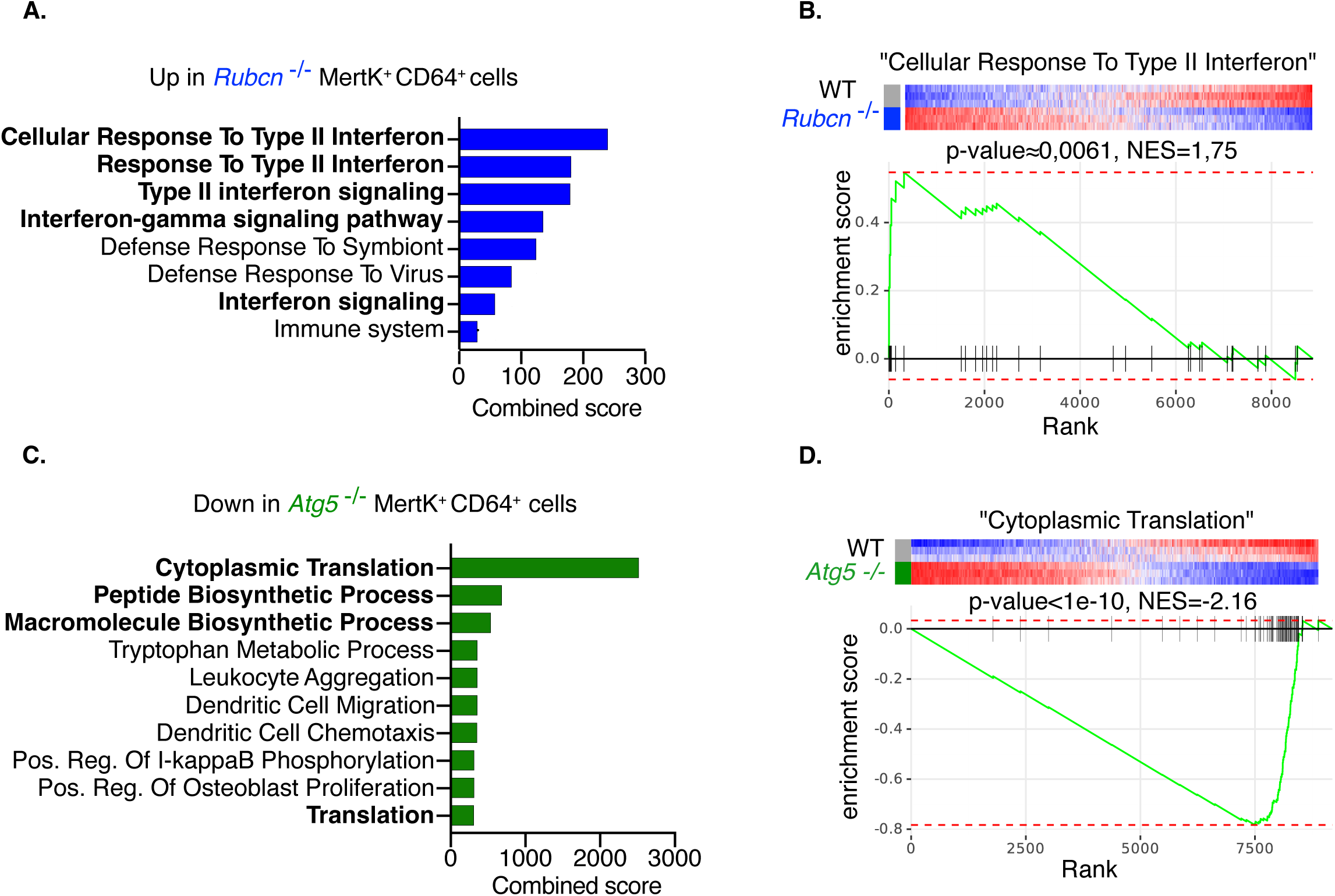
Pathways affected by autophagy modulation in Mertk+ FcyRI+ splenic phagocytes. (**A**) Pathways upregulated in Rubicon deficient (Rubcn -/-) Mertk+ FcyRI+ splenic phagocytes. (**B**) GSEA of cellular response to type II interferon in Rubcn -/- and wild-type Mertk+ FcyRI+ phagocytic cells. (**C**) Pathways downregulated in Atg5 -/- Mertk+ FcyRI+ phagocytic cells. (**D**) GSEA of cytoplasmic translation in Atg5 -/- and wild-type Mertk+ FcyRI+ splenic phagocytes.

**Supplementary Figure 5:**
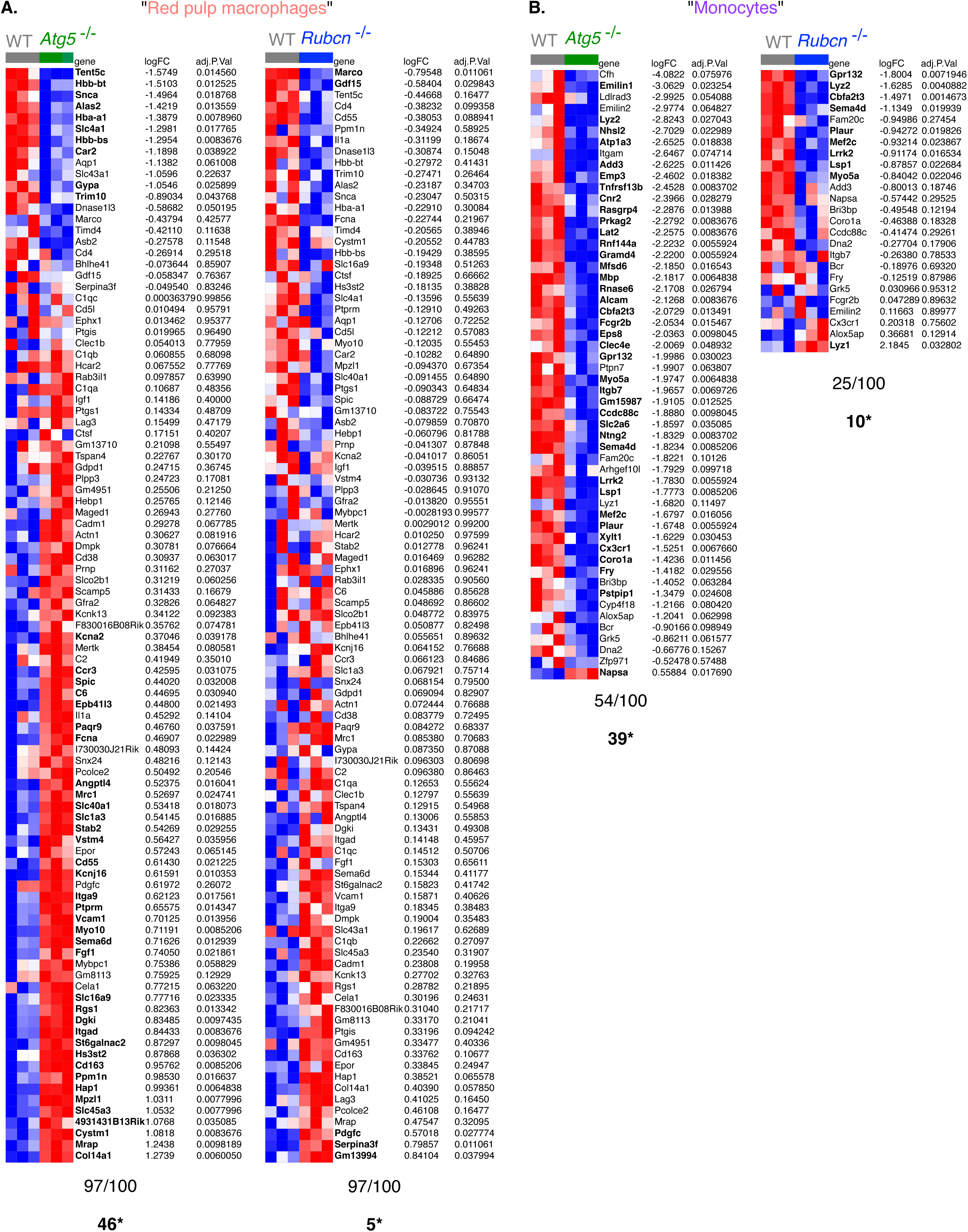
Modulation of autophagy significantly affects the monocyte but not the macrophage population in the spleen. (**A**) Heat maps showing genes from the red pulp macrophage signature in Atg5 -/-, Rubcn -/- Mertk+ FcyRI+ splenic phagocytes and their wild-type counterparts. The number of genes detected in each dataset and the number of significantly enriched genes are shown below. (**B**) Heat maps showing genes from the "monocyte" signature in Atg5 -/-, Rubcn -/- Mertk+ FcyRI+ splenic phagocytes and their wild-type counterparts. The number of genes detected in each dataset and the number of significantly enriched genes are shown below.

**Supplementary Figure 6:**
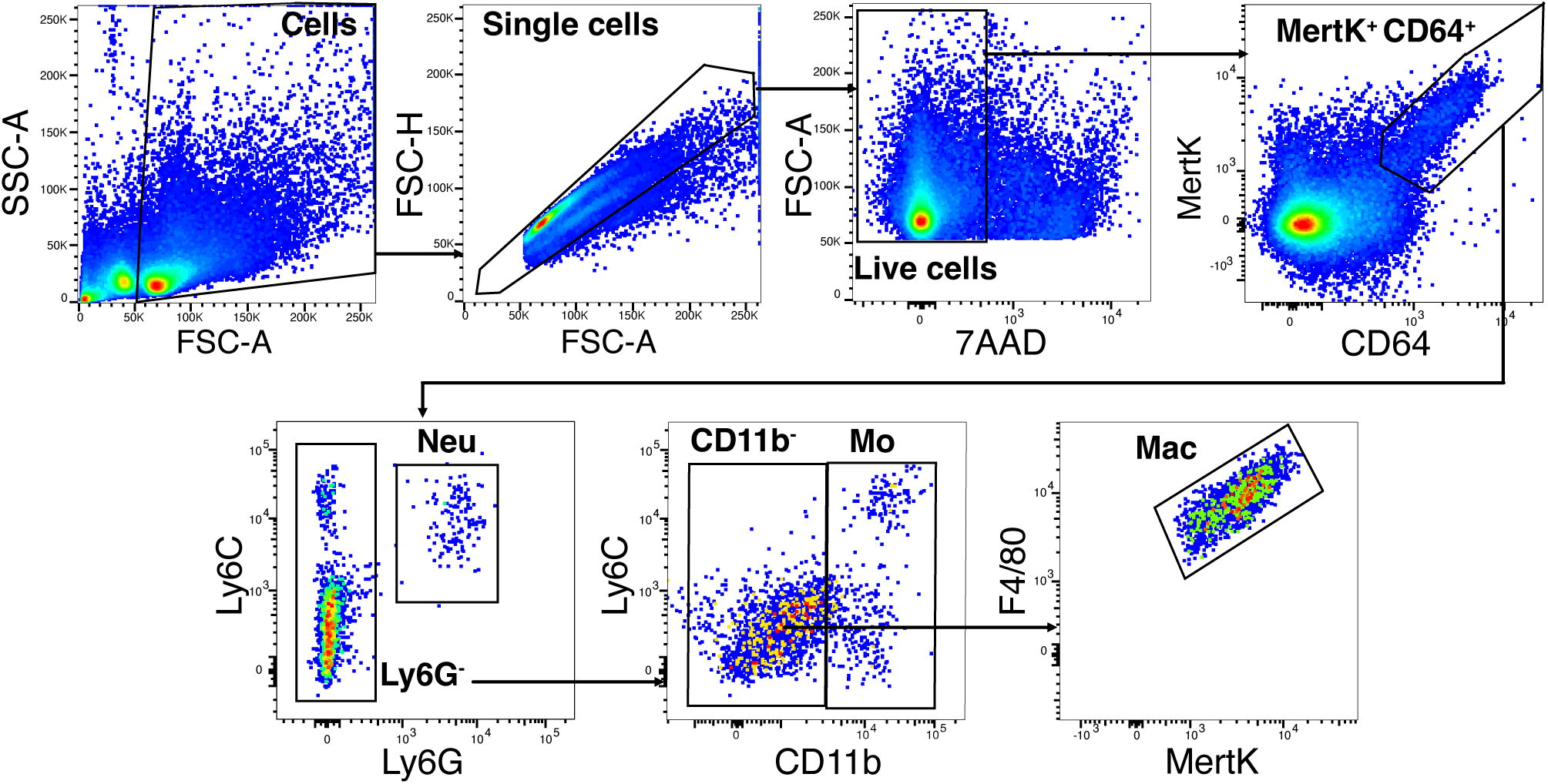
Mertk+ FcyRI+ splenic phagocytes contain mainly monocytes and red pulp macrophages. (**A**) Gating strategy used to analyze other myeloid cell populations within the Mertk+ FcyRI+ splenocyte gate.

**Supplementary Figure 7:**
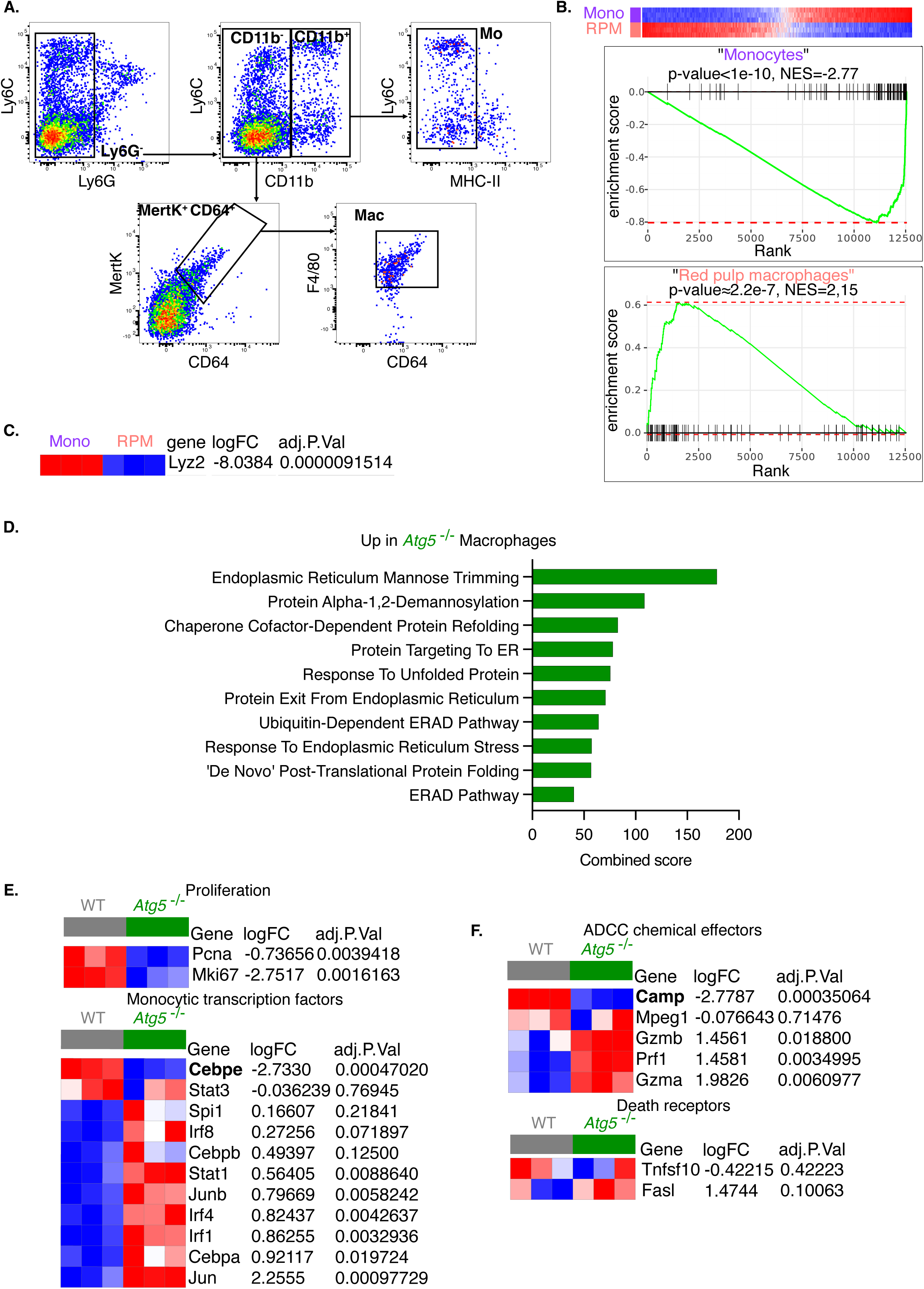
Monocytes and red pulp macrophages are differentially affected by Atg5-/- deletion. (**A**) Gating strategy used to sort monocytes and red pulp macrophages from spleens of Atg5-/- and wild-type mice. (**B**) GSEA of monocyte and red pulp macrophage signatures in monocytes and red pulp macrophages from the spleen of wild-type mice. Phantasus was used to visualize the data. (**C**) Lyz2 expression in monocytes and red pulp macrophages from the spleen of wild-type mice. (**D**) Pathways upregulated in Atg5-/- red pulp macrophages. (**E-F**) Heat maps showing genes involved in cell proliferation, monocyte differentiation and effectors for ADCC in monocytes from spleen of Atg5 -/- or wild-type mice.

## Data Availability

All data supporting the conclusions are presented in the manuscript and the supplementary materials (Supplementary Figures, source data files).

## Acknowledgements

This work was supported by grants from the “Agence Nationale de la Recherche” (ANR-11-IDEX-0005-02 Laboratory of Excellence INFLAMEX to LS), the CSL Behring Foundation, the “Fondation pour la Recherche Medicale” (fellowship to DK), the World Wide Cancer Reasearch (fellowship to DK) and La Ligue Contre le Cancer (PhD fellowship to MN).

